# Azadiradione regulates Heat Shock Factor 1 function by interacting with its DNA-binding domain independent of the oligomerization domain

**DOI:** 10.1101/2025.04.28.650926

**Authors:** Anirban Manna, Papri Basak, Chirantan Majumder, Nilanjana Hazra, Kuladip Jana, Jayanta Mukhopadhyay, Subhash C. Mandal, Mahadeb Pal

## Abstract

Heat shock factor 1 (HSF1) masters cellular proteostasis under stress by upregulating the expression of molecular chaperones that help refold or degrade the misfolded proteins. HSF1 activation involves a monomer-to-oligomer transition and binding to its recognition sequence, the heat shock elements (HSEs) on its target gene promoters. HSF1 activity declines with age as well as in neurodegenerative disorders (NDs) such as Parkinson’s disease, highlighting the need for strategies to restore its function. Azadiradione (AZD), a limonoid isolated from *Azadirachta indica* seeds, directly activates HSF1 in cellular and preclinical ND models, unlike other small-molecule activators reported elsewhere. We investigated the molecular basis of AZD-mediated HSF1 activation using purified variants of this protein including those without its oligomerization and transactivation domain. Fluorescence polarization and dynamic light scattering assays revealed that AZD promotes the oligomerization of monomeric HSF1 to enhance its HSE-binding affinity by engaging with its DNA-binding domain (DBD). The oligomerization domain known to be required for stress-induced activation, appears redundant in AZD-mediated activation. Furthermore, evidence suggests AZD-induced conformational alterations in the HSE facilitate its binding to the HSF1 monomer. Notably, AZD reduces the DNA-binding ability of pre-assembled oligomeric HSF1 by triggering its amyloid-like aggregation. This finding also highlighted a potential anticancer effect of AZD, as cancer cells heavily depend on this HSF1 population for rapid proliferation and survival. Overall, these findings offer novel insights into the functional regulation of HSF1 and suggest a framework for developing small-molecule HSF1 activators with therapeutic potential for protein conformation disorders.

## Introduction

Cells initiate the heat shock response (HSR), an evolutionarily conserved transcriptional program to maintain homeostasis under stressful conditions [1]. Heat shock factor 1 (HSF1), the master regulator of HSR in eukaryotic cells, mediates this process by upregulating its target genes’ expressions, which includes the inducible molecular chaperones. They help refold, degrade, and clear the misfolded proteins to mitigate the toxicity induced by cellular stress [2,3]. HSF1 binds its DNA recognition motif called the heat shock element (HSE), consisting of contiguous inverted repeats of 5’-nGAAn-3’ sequence on the target genes’ promoters as a homotrimer/oligomer to upregulate their expressions [4].

Human HSF1 (MW: ∼57 kDa) is a multidomain protein. Its N-terminal DNA-binding domain (DBD) facilitates the binding of oligomeric HSF1 to the HSE. The highly conserved DBD is the only structurally characterized domain of HSF1. This domain is followed by an α-helical leucine-rich oligomerization domain comprising three leucine zippers (LZ1–3), which mediates stress-induced multimerization of monomeric HSF1. An intrinsically disordered regulatory domain (RD), located C-terminal to the LZ1–3 domain, represses HSF1 activity by undergoing post-translational modifications. Another short leucine zipper domain (LZ4) lies between the RD and the C-terminal transactivation domain (TAD). The partly α-helical and partly unstructured TAD is believed to interact with various transcriptional cofactors [5].

Under normal conditions, HSF1 remains inactive due to intramolecular interactions between its LZ4 and LZ1–3 domains. Thermal shock triggers the unfolding of LZ4 (and possibly LZ1–3), resulting in their dissociation. This allows LZ1-3-mediated intermolecular coiled-coil interactions among individual HSF1 protomers, leading to the homo-trimerization/ - oligomerization of monomeric HSF1 [6].

Individuals with neurodegenerative diseases (NDs) exhibit a buildup of toxic protein aggregates in various regions of the central nervous system, ultimately leading to neuronal death [7,8]. Notably, the HSR is impaired in ND patients [7,9]. Thus, forcibly upregulating or activating HSF1 is expected to aid the clearance of toxic protein aggregates, as demonstrated in fruit fly and mouse models of Huntington’s disease [10–12].

In contrast to normal cells, which deactivate HSF1 after stress resolution, cancer cells maintain HSF1 in a constitutively activated state [13,14]. Cancer cells require elevated levels of molecular chaperone to sustain their proteome and proliferative potential. Additionally, cancer cells rewire the HSF1 transcriptome to support their needs, targeting genes involved in critical cellular functions—a phenomenon termed the ‘HSF1 cancer program’ [15]. Therefore, pharmacological inhibition of HSF1 represents a promising anticancer strategy [16].

Several small-molecule activators of HSF1 are reported in the literature that were shown to enhance HSF1 activity indirectly, primarily by inhibiting the cellular Hsp90 or the proteasome [8,17–19]. Nelson and colleagues (2016) identified azadiradione (AZD, MW: 451 Da), a small molecule purified from *Azadirachta indica* (Neem) seeds, as a direct activator of HSF1. AZD induces HSF1 homo-multimerization, thereby increasing its affinity for HSE without affecting cellular Hsp90, the proteasome, or ROS levels. To date, AZD remains the only reported direct activator of HSF1. It has been shown to clear toxic polyglutamine protein aggregates in fruit fly and mouse models of NDs [12,20]. Surprisingly, AZD activates HSF1 and upregulates molecular chaperone expression even in the absence of proteotoxic stressors at nontoxic cellular concentrations [20]. Furthermore, AZD could inhibit Tau protein aggregation and reduce Tau aggregate-induced toxicity in HEK293T cells at low concentrations [21].

The present study investigates the underlying molecular basis of AZD-mediated modulation of HSF1 activity to gain deeper insights into the HSR and develop better HSF1 modulators in the future. We aimed to determine the binding site(s) of AZD on human HSF1 and the mechanism(s) involved in AZD-induced activation of HSF1 in the absence of proteotoxic stress. We employed various biochemical and biophysical tools to study the interactions of HSF1 and its variants with AZD *in vitro*. We demonstrate that AZD induces the oligomerization of monomeric HSF1 to increase its HSE-binding affinity by interacting with the DBD. The oligomerization domain, indispensable for stress-induced HSF1 activation, appears non-essential for AZD-induced oligomerization and HSE-binding. We further show that AZD-induced conformational changes in the HSE facilitate the interaction of monomeric HSF1 with HSE. Surprisingly, AZD could induce the destabilization/ aggregation of the preassembled oligomeric forms of HSF1, compromising their HSE-binding activities. Cancer cells are known to be excessively dependent on oligomeric HSF1 for their survival and proliferation, underscoring the compound’s potential use as an anticancer agent.

## Results

### Expression and purification of monomeric and oligomeric species of HSF1

HSF1-WT and its variants were overexpressed in *E. coli* BL21(λDE3) to analyze their interactions with AZD (**Fig. 1A-B**). HSF1-WT isolated from the overexpressing *E. coli* strain by affinity chromatography occurs as a mixture of multiple molecular species [4,6]. The monomeric and oligomeric molecular species of HSF1 were separated by size exclusion chromatography (SEC) as described in ‘Materials & Methods’ **(Fig. 1 C-E).** The SEC profile of HSF1-WT revealed two prominent peaks: a taller peak (peak 1, red arrow) eluting at 8.77 ml and a shorter one at 12.44 ml (peak 3, purple arrow). In addition, a small hump eluting at 10.59 ml was denoted as peak 2 **(Fig. 1C).** HSF1-ΔLZ1-3, constitutively monomeric (CM), shows a major peak eluting at 11.86 ml (peak 1’, purple arrow), close to the elution volume (EV) of peak 3 of HSF1-WT **(Fig. 1D).** On the other hand, HSF1-LZ4m, constitutively trimeric (CT), shows a major peak at 8.52 ml (peak 1”, red arrow), close to the EV of peak 1 of HSF1-WT **(Fig. 1E).** As expected, a little faster electrophoretic mobility of HSF1-ΔLZ1-3 (peak 1’) relative to HSF1-WT (peak 3) was revealed in a blue native gel. Peak 1 of HSF1-WT (8.77 ml) and peak 1” of HSF1-LZ4m (8.52 ml) also showed very similar electrophoretic mobilities **(Fig. 1F).**

**Figure 1:**
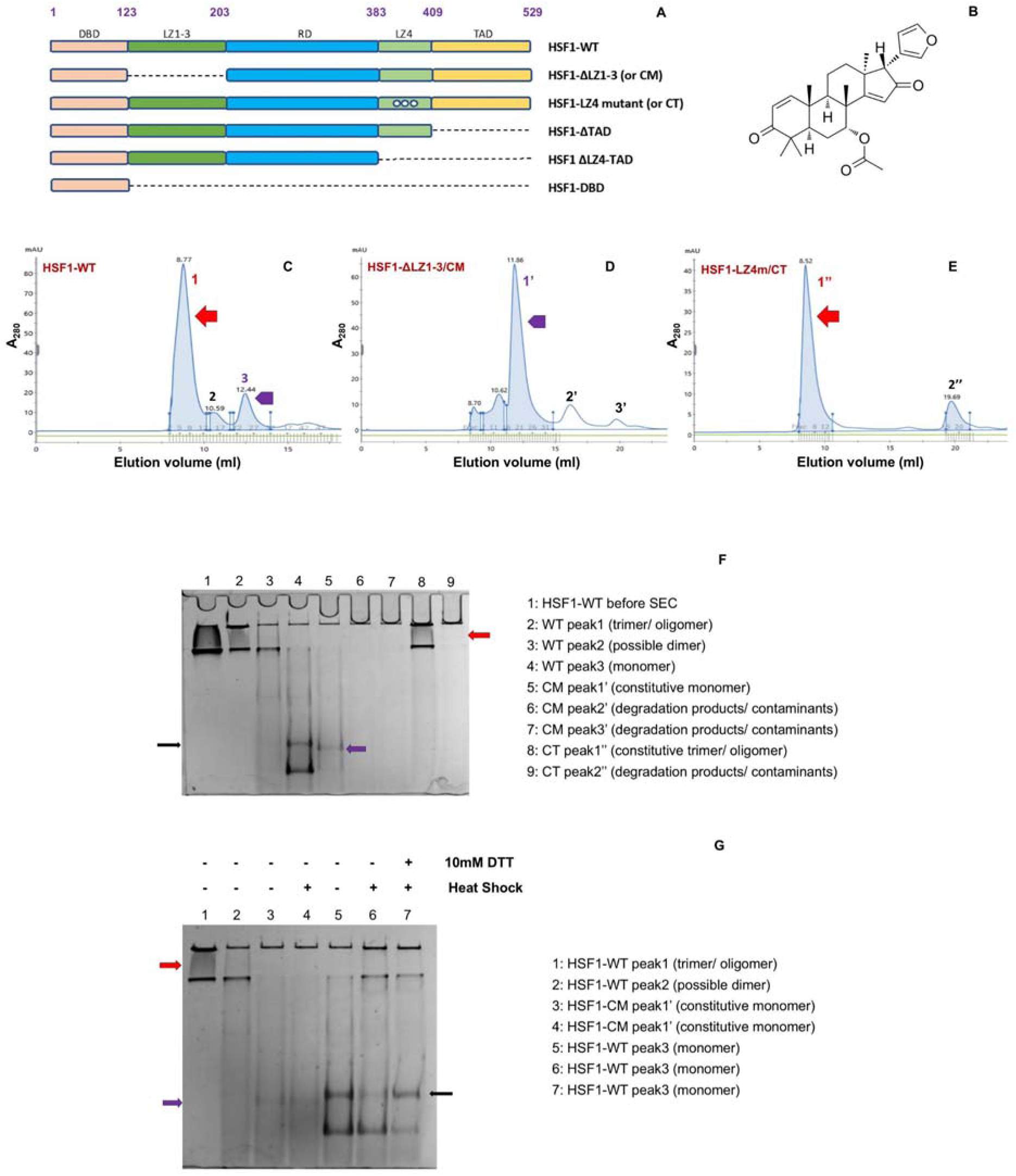
Isolation of monomeric and oligomeric species of human HSF1. A: The domain organization of full-length human HSF1 and its derivatives used in the present study. Amino acid residues marking the domain boundaries are indicated in purple. [CM: Constitutive Monomer; CT: Constitutive Trimer; TAD: Transactivation Domain; DBD: DNA-Binding Domain; LZ: Leucine Zipper] **B.** Molecular structure of Azadiradione (AZD) C-E. Size exclusion chromatography (SEC) profiles of affinity-purified HSF1 and its derivatives. Numerical representation denotes the distinct peaks of different molecular species of HSF1 and its variants. **(C)** Red arrow: trimeric/ oligomeric species of HSF1-WT; purple arrow: monomeric species of HSF1-WT **(D)** Purple arrow: HSF1-constitutive monomer (CM) **(E)** Red arrow: HSF1-constitutive trimer/ oligomer (CT) **F.** Resolution of the molecular species of HSF1 and its derivatives obtained by SEC (shown in C-E) using Blue Native PAGE (7%). The black arrow indicates the HSF1-WT monomer, the purple arrow indicates the HSF1-CM and the red arrow indicates the trimeric/ oligomeric species of both HSF1-WT and -CT. **G.** Confirmation of SEC-purified HSF1-WT monomer by Blue Native PAGE. The black arrow indicates the HSF1-WT monomer. HSF1-CM is indicated by the purple arrow and HSF1-WT trimer/ oligomer by the red arrow.

**Table 1:**
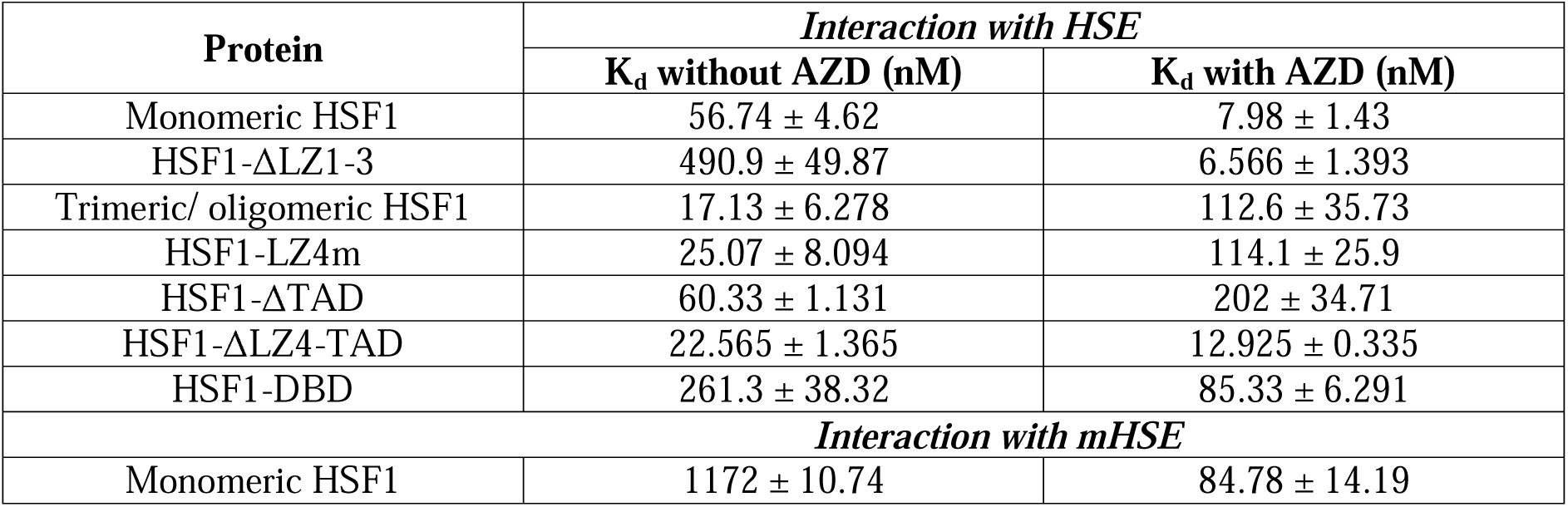
Results of the protein-DNA interaction studies performed by fluorescence polarization assay

The heat shock (HS) treatment of HSF1-WT (peak 3) led to its oligomerization, visible as a band shift to a higher MW position on a blue native gel. DTT partially reversed this shift, restoring the original band position corresponding to the non-heat-shocked sample[30]. In contrast, HS did not alter the band position of peak 1’ of HSF1-ΔLZ1-3, a mutant defective in oligomerization **(Fig. 1G).** This indicates that peak 3 of HSF1-WT represents the monomeric species.

### Monomeric HSF1 forms oligomers upon incubation with AZD and binds the HSE with enhanced affinity

To investigate the effect of purified AZD **(Fig. 1B)** on the HSF1-HSE binding affinity, we analysed the binding of wild-type HSF1 and its variants with a 5’FAM-tagged HSE (mentioned in ‘Materials and Methods’) through fluorescence polarization (FP) assay. In the absence of protein, the HSE exhibits high rotational diffusion, resulting in low FP values, while its binding with the protein restricts this rotational diffusion, increasing the polarization values. By measuring the increase in polarization during the titration of protein into a solution containing a fixed concentration of HSE, a binding curve was generated. This curve was used to determine the equilibrium dissociation constant (K_d_) of the protein-HSE interaction.

The interaction of monomeric HSF1 with HSE in the absence of AZD yielded a K_d_ value of 56.74 ± 4.62 nM, which dramatically reduced to 7.98 ± 1.43 nM in the presence of 10 μM AZD, indicating an AZD-induced increase in the binding affinity of monomeric HSF1 with HSE by ∼7.1-fold **(Fig. 2: A-C).** To understand if this increase in HSE-binding affinity was due to the formation of trimeric/ higher oligomeric structures, we performed a dynamic light scattering (DLS) assay. An addition of AZD to monomeric HSF1 appreciably increased the hydrodynamic radius of the protein independent of HSE **(Fig. 2G).** Therefore, AZD induces the homo-multimerization of HSF1-WT monomer in the absence of HSE under stress-free conditions, which increases its binding affinity for the HSE.

**Figure 2:**
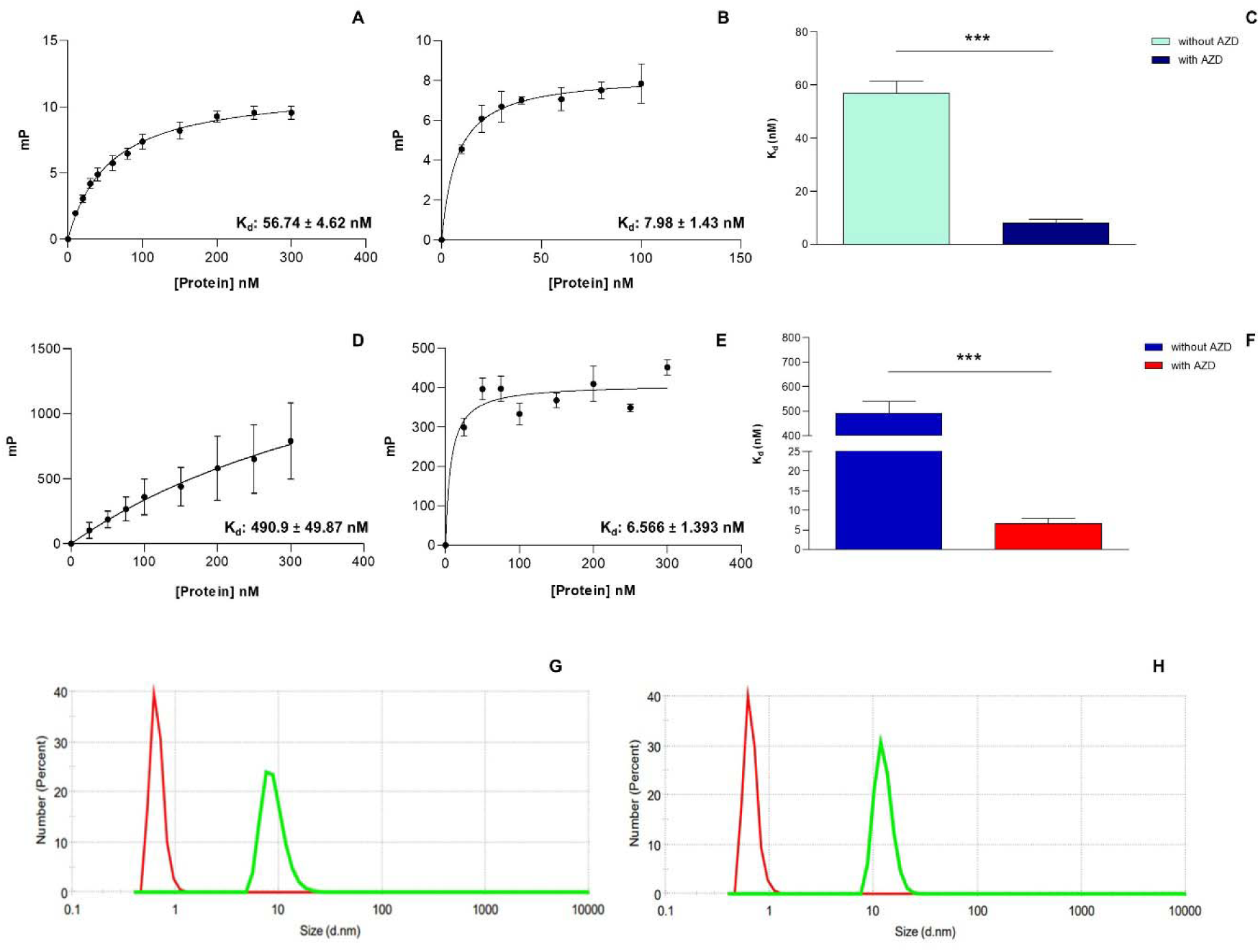
AZD increases the binding affinity of both HSF1-WT monomer and HSF1-CM for FAM-labelled HSE (FAM-HSE) by inducing a monomer to multimer transition, as determined by Fluorescence Polarization (FP) and Dynamic Light Scattering (DLS) assays. Dissociation constants (K_d_) obtained by plotting the millipolarization value of FAM-HSE against increasing concentrations of HSF1-WT monomer, in the absence of AZD (A) and in the presence of 10 μM AZD (B); of HSF1-CM, in the absence of AZD (D) and in the presence of 10 μM AZD (E). The bar graphs C and F compare the K_d_ values of the interactions obtained for HSF1-WT monomer (A, B) and HSF1-CM (D, E), respectively. For HSF1-WT monomer, n=3 and p-value is 0.0005 (***). For HSF1-CM, n=4 and p-value <0.0001 (***). DLS assays of AZD-induced oligomerization of HSF1-WT monomer (G) and HSF1-CM (H), in the absence of HSE. The red and green peaks correspond to the protein size distribution without and with AZD, respectively.

### AZD increases the binding affinity of the constitutively monomeric HSF1 variant for the HSE

The oligomerization domain of HSF1 (LZ1-3) is implicated in the HS-induced trimerization/ oligomerization of the protein through the formation of an inter-subunit triple-stranded coiled-coil structure[4]. A construct lacking this domain (HSF1-ΔLZ1-3 or HSF1-CM) did not oligomerize upon heat shock, as the individual subunits could not be held together in a complex in the absence of the leucine zipper **(Fig. 1 A, G).** We analyzed the effect of AZD on this construct and compared it with that on wild-type monomeric HSF1. As expected, HSF1-ΔLZ1-3 bound HSE with a very low affinity (K_d_: 490.9 ± 49.87 nM). However, to our astonishment, the presence of AZD increased the binding affinity by ∼75 fold (K_d_: 6.566 ± 1.393 nM) **(Fig. 2: D-F).** Thus, consistent with earlier reports, HSF1-ΔLZ1-3 binds HSE with a much lower affinity compared to the HSF1-WT monomer [4]. Intriguingly, the affinity of HSF1-CM-HSE interaction becomes comparable to that of HSF1-WT monomer-HSE interaction in the presence of AZD. Congruently, in our DLS assay, the hydrodynamic radius of HSF1-ΔLZ1-3 increased considerably upon incubation with AZD independent of HSE, like the HSF1-WT monomer **(Fig. 2H).** Therefore, AZD-induced multimerization and DNA binding of monomeric HSF1 seem independent of its oligomerization domain, in stark contrast to HS-induced HSF1 activation.

### AZD interrupts the binding of trimeric/oligomeric HSF1 with the HSE

Monomeric HSF1 has been demonstrated to undergo trimerization/oligomerization upon stress exposure to bind the HSE in its target gene promoters with high affinity [17]. We wished to study the effect of AZD on the interaction of the pre-assembled trimeric/oligomeric form of HSF1-WT (isolated by SEC) with the HSE **(Fig. 1C).** Surprisingly, AZD weakened the binding of this species with HSE by ∼6.57-fold (K_d_: 112.6 ± 35.73 nM with AZD vs 17.13 ± 6.278 nM without AZD) **(Fig. 3: A-C).** Notably, AZD increased the binding affinity of HSF1-WT monomer with HSE by ∼7.1-fold **(Fig. 2: A-C).**

**Figure 3:**
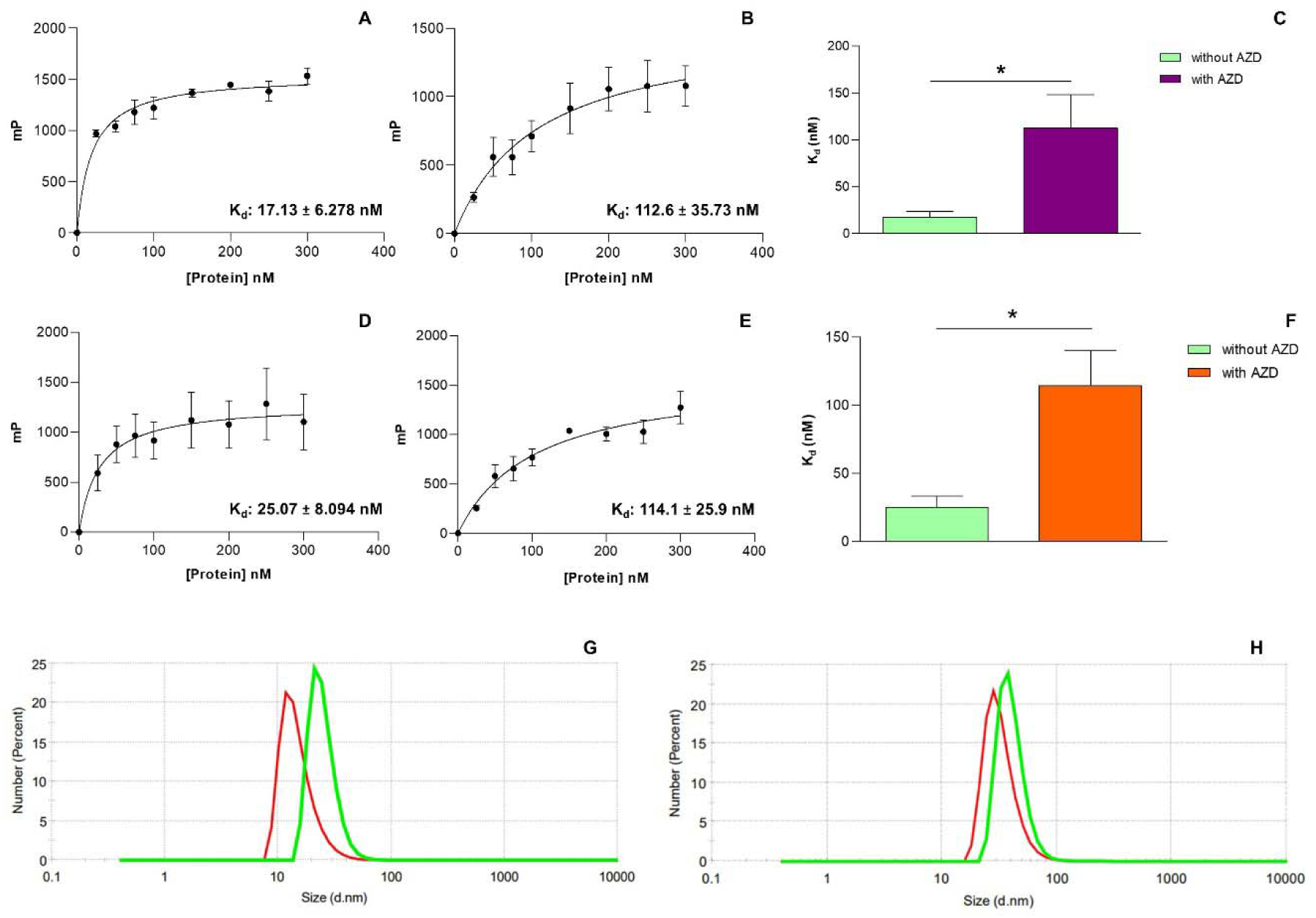
AZD disrupts the interaction of HSF1-WT trimer and HSF1-CT with FAM-HSE by inducing structural destabilization of these proteins, as demonstrated by FP and DLS assays. Dissociation constants (K_d_) obtained by plotting the millipolarization value of FAM-HSE against increasing concentrations of HSF1-WT trimer, in the absence of AZD (A) and in the presence of 10 μM AZD (B); of HSF1-CT, in the absence of AZD (D) and in the presence of 10 μM AZD (E). The bar graphs C and F compare the K_d_ values of the interactions obtained for HSF1-WT trimer (A, B) and HSF1-CT (D, E), respectively. For the HSF1-WT trimer, n=3 and p-value is 0.0269 (*). For HSF1-CT, n=4 and p-value is 0.0168 (*). DLS assays depict the oligomeric status of HSF1-WT trimer (G) and HSF1-CT (H), in the absence of HSE. The red and green peaks correspond to the protein size distribution without and with AZD, respectively.

Next, to better understand the interaction of AZD with trimeric/ oligomeric HSF1, we analyzed the effect of AZD on the interaction of HSF1-LZ4m with the HSE. As mentioned earlier, this mutant remains constitutively trimeric/ oligomeric, exhibiting the same SEC elution profile as the HSF1-WT trimer/ oligomer **(Fig. 1: C & E).** Like trimeric/ oligomeric HSF1-WT, AZD weakened the interaction of this species with HSE by ∼4.5-fold (K_d_: 25.07 ± 8.094 nM without AZD vs 114.1 ± 25.9 nM with AZD). **(Fig. 3: D-F).**

To check whether this decrease in HSE-binding affinity is caused by an AZD-induced dissociation of the oligomeric structure (oligomer-to-monomer transition), we performed DLS analyses. Interestingly, DLS studies revealed a small increase in the hydrodynamic radii of these two forms of HSF1 upon incubation with AZD **(Fig. 3 G & H),** suggesting an AZD-induced destabilization of the structure of the proteins, accounting for their weakened binding with HSE [31].

### The DNA-binding domain (DBD) of HSF1 plays a critical role in the AZD-induced binding of monomeric HSF1 to the HSE

We tested the C-terminal truncation derivatives of HSF1 to gain insights into the domain(s) involved in AZD-induced binding of HSF1 to HSE **(Fig. 1A).** HSF1-ΔTAD was shown to bind HSE with a K_d_ of 60.33 ± 1.131 nM, which was increased to 202 ± 34.71 nM in the presence of AZD. DLS study showed a small increase in the hydrodynamic radius of HSF1-ΔTAD upon incubation with AZD in the absence of HSE **(Fig. 4: A, B, C & J).** This result indicates a destabilization of the protein’s structure upon AZD binding, causing the reduction in HSE-binding affinity.

**Figure 4:**
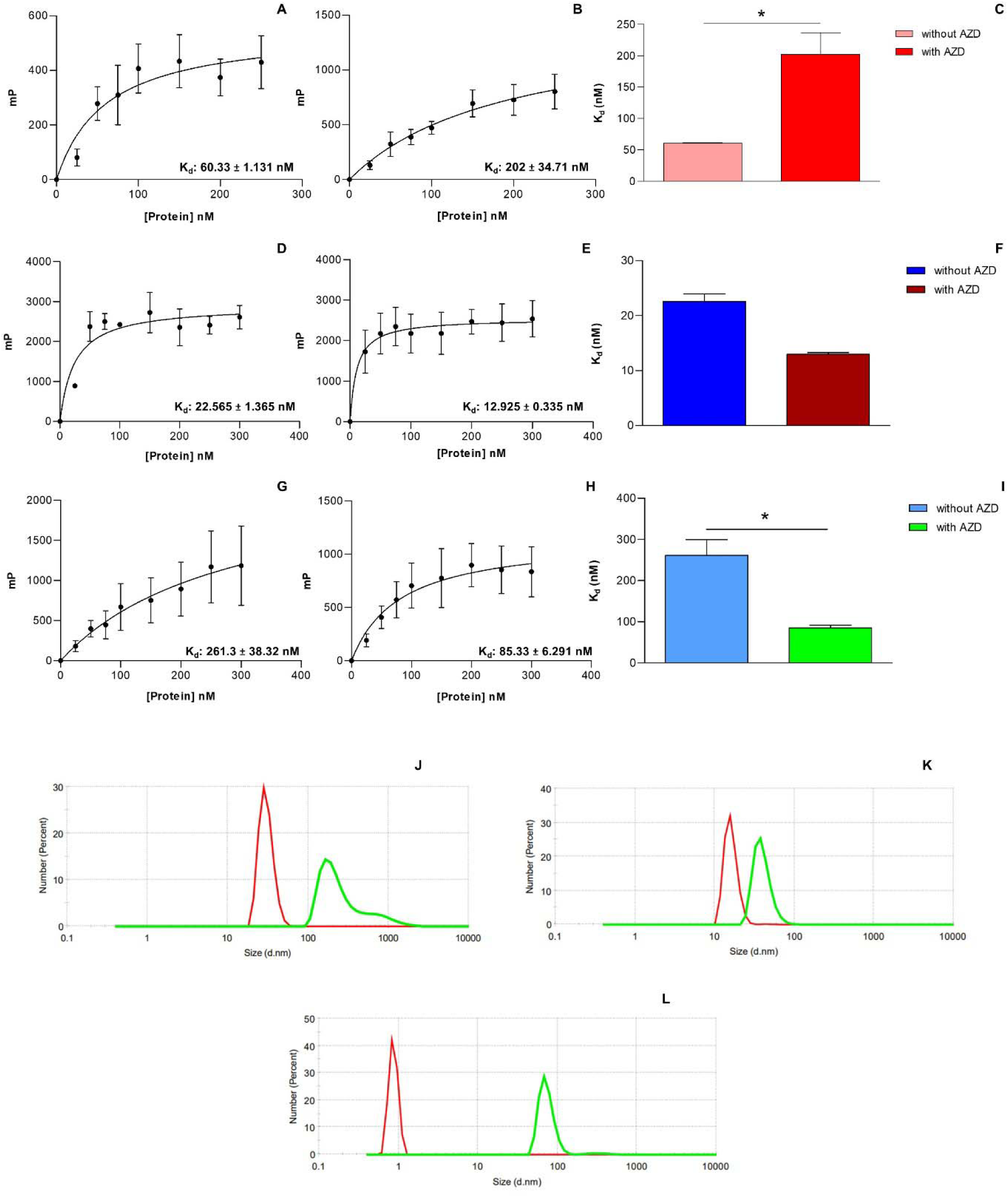
AZD exerts varied effects on the HSE-binding ability of different C-terminal truncation derivatives of HSF1, as demonstrated by FP and DLS assays. Dissociation constant (K_d_) obtained by plotting the millipolarization values of FAM-HSE against increasing concentrations of HSF1-ΔTAD in the absence (A) and presence of 10 μM AZD (B); of HSF1-ΔLZ4-TAD in the absence (D) and presence of 10 μM AZD (E); of HSF1-DBD in the absence (G) and presence of 10 μM AZD (H). The bar graphs C, F and I compare the K_d_ values of the interactions obtained for HSF1-ΔTAD (A, B), HSF1-ΔLZ4-TAD (D, E), and HSF1-DBD (G, H), respectively. For HSF1-ΔTAD, n=3 and p-value is 0.0151 (*). For HSF1-ΔLZ4-TAD, n=2. For HSF1-DBD, n=3 and p-value is 0.0106 (*). DLS assays depict the oligomeric status of HSF1- ΔTAD (J), HSF1- ΔLZ4-TAD (K), and HSF1-DBD (L) in the absence of HSE. The red and green peaks correspond to the protein size distribution without and with AZD, respectively.

HSF1-ΔLZ4-TAD (consisting of the DBD, LZ1-3 and RD) is expected to remain an oligomer, as it lacks the autoinhibitory LZ4 domain. This protein bound HSE with a K_d_ of 22.56 ± 1.365 nM in the absence of AZD. The binding affinity increased by a small margin (K_d_: 12.92 ± 0.335 nM) in the presence of AZD. DLS study indicated a slight increase in the protein’s hydrodynamic radius upon AZD exposure. We did not analyze this protein further owing to its relative insensitivity to AZD **(Fig. D, E, F& K).**

The DBD of HSF1 enables the protein to bind the HSE **(Fig. 1A).** The binding affinity of purified HSF1-DBD with HSE is expected to be low, owing to the absence of the oligomerization domain. Although certain residues have been shown to mediate DBD-DBD contacts in the crystal structure of the DBD-HSE complex, these seem insufficient for maintaining a stable trimeric/oligomeric structure [32]. Indeed, HSF1-DBD was found to bind HSE with a low affinity (K_d_: 261.3 ± 38.32 nM). The presence of AZD, however, increased the binding affinity threefold (K_d_: 85.33 ± 6.291 nM). The AZD-induced increase in the hydrodynamic radius of HSF1-DBD shown by our DLS study suggested the formation of a multimeric structure upon incubation of the protein with AZD, independent of HSE **(Fig. 4: G, H, I & L).**

### AZD interacts with the monomeric forms of HSF1 in the absence of HSE

The interaction of AZD with HSF1-WT monomer, HSF1-CM, and HSF1-DBD caused multimerization and stabilization of these proteins and increased their binding affinities for HSE, as indicated by our FP and DLS analyses. Therefore, we were interested to know the binding affinities of these proteins for AZD in the absence of HSE.

For this purpose, intrinsic fluorescence spectra of AZD-equilibrated HSF1-WT monomer, CM, and DBD were individually recorded as described in ‘Materials and Methods.’ An AZD-induced enhancement in the intrinsic fluorescence intensity of all the proteins was observed upon the excitation of their tryptophan and tyrosine residues at 280 nm. The fluorescence intensity of all the proteins increased considerably at lower AZD concentrations before reaching saturation in fluorescence enhancement **(Fig. 5: A-F).** The K_d_ for all the protein-ligand interactions were quite close. AZD in buffer showed negligible fluorescence emission under the conditions tested.

**Figure 5:**
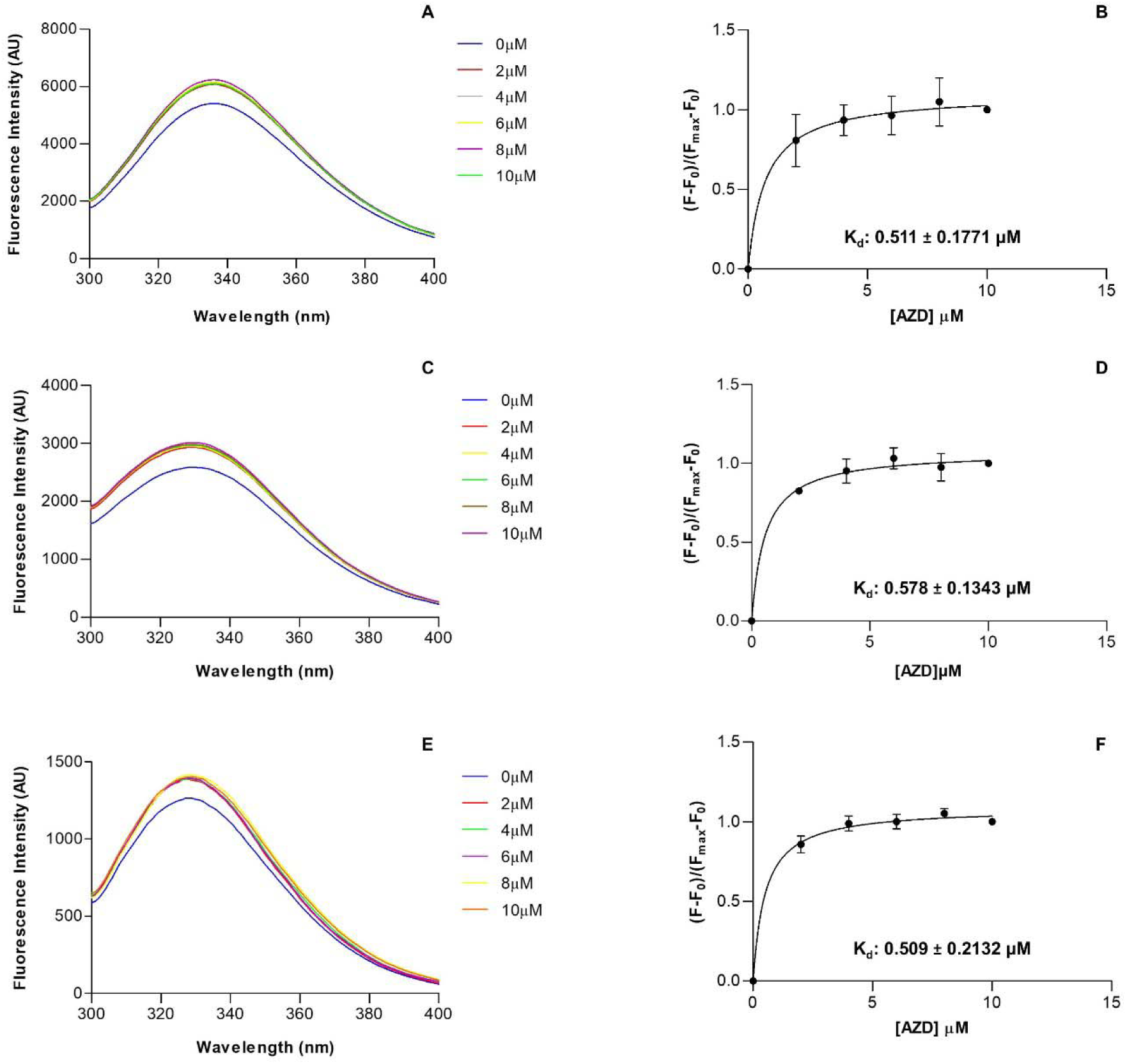
AZD interacts with the monomeric forms of HSF1 (in the absence of HSE) with comparable binding affinities as studied by protein intrinsic fluorescence spectroscopy. Intrinsic fluorescence emission spectra of HSF1-WT monomer (A), HSF1-CM (C), and HSF1- DBD (E), obtained after incubation with increasing concentrations of AZD, as indicated. Dissociation constant (K_d_) obtained from the binding curves of 2 μM HSF1-WT monomer (B), HSF1-CM (D), and HSF1-DBD (F) with increasing concentrations of AZD, as indicated (n=3 for each protein). For each concentration of AZD, the fluorescence emission intensity of HSF1-WT monomer at 335 nm, HSF1-CM at 330 nm, and HSF1-DBD at 328 nm (λ_max_) were considered for plotting the curves.

### AZD interacts with HSE and mutant HSE in the absence of protein

**Nelson *et al.* (2016)** analysed the possible interaction(s) of AZD with HSE and HSF1-DBD *in silico* by docking AZD in the crystal structure published by **Neudegger *et al.* (2016),** which indicated the binding of AZD at the protein-DNA interface, contacting both the HSE and HSF1- DBD [20,32]. In the crystal structure, however, a two-site HSE was used [32]. Here, we analysed the interaction of canonical three-site HSE with AZD *in vitro* in the absence of protein by fluorescence spectroscopy. Upon titration of FAM-labelled HSE (10 nM) with increasing concentrations of AZD, the fluorescence intensity at 520 nm was found to increase in a concentration-dependent manner, ultimately reaching saturation (λ_ex_: 495 nm). The fluorescence of AZD in the buffer was negligible in the range tested. The K_d_ of this interaction was 1.315 ± 0.1266 μM. This suggested an interaction of AZD with the DNA molecule, occupying specific binding site(s). Gradually increasing the concentration of AZD saturated these site(s) **(Fig. 6: A & B).**

**Figure 6:**
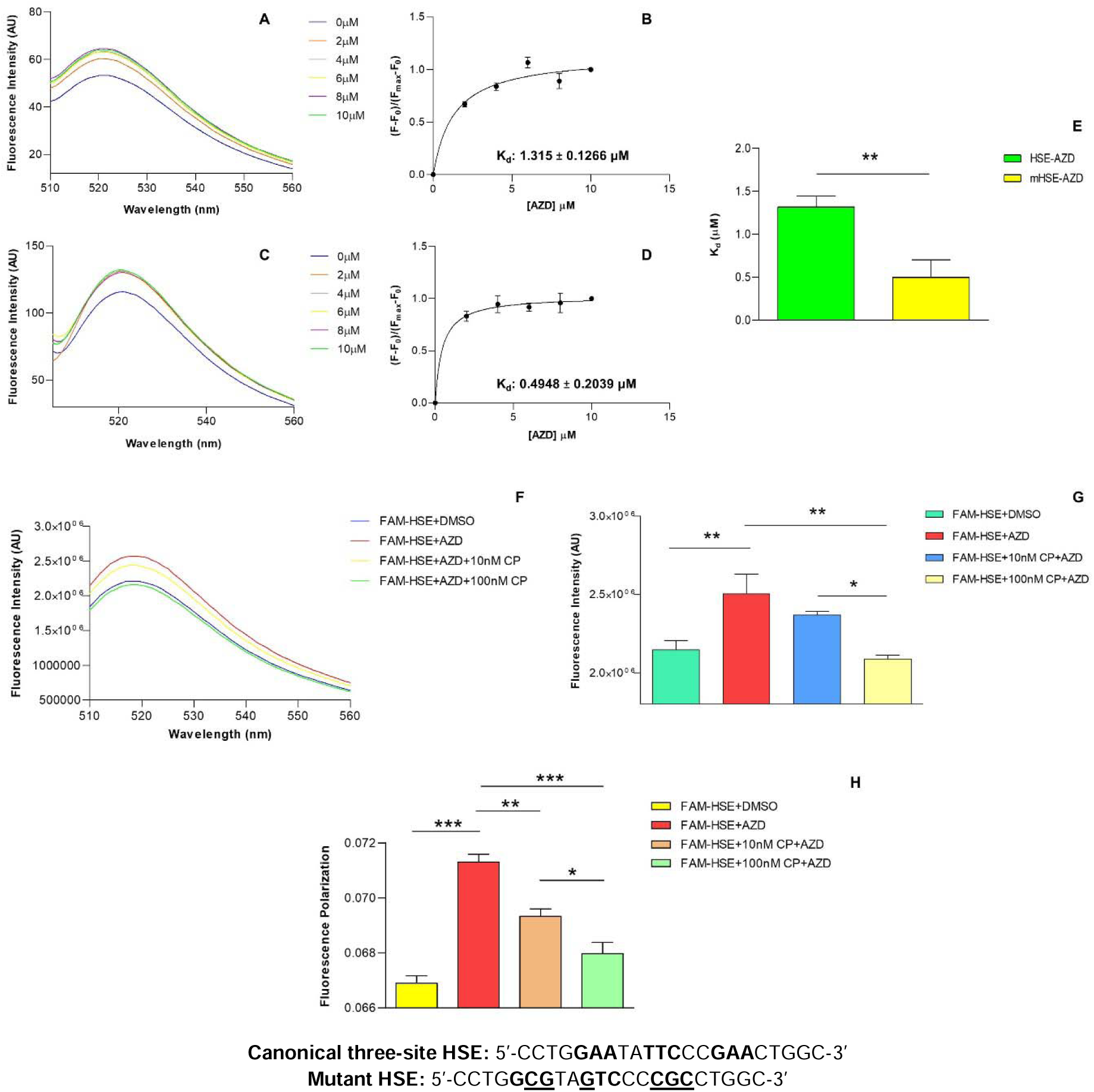
AZD interacts with canonical and mutant HSE-DNA in the absence of protein. Fluorescence emission spectra of FAM-labelled HSE (A) and mHSE (C) upon incubation with increasing concentrations of AZD, as indicated. Dissociation constants (K_d_) obtained from the binding curves of 10 nM FAM-labelled HSE (B) and mHSE (D) with increasing concentrations of AZD, as indicated. For each concentration of AZD, the fluorescence emission intensity at 520 nm (λ_max_) was considered for plotting the curves. (E) Bar graph comparing the K_d_ for the HSE-AZD and mHSE-AZD interactions [n=4 and p- value is 0.009 (**)]. (F) Fluorescence emission spectrum of FAM-labelled HSE-AZD mixture upon incubation with unlabelled HSE (Cold Probe; CP). Two different concentrations of CP were used in this experiment, as indicated. (G) Bar graph depicting the change in fluorescence emission intensity of FAM-labelled HSE at 520 nm upon incubation with AZD in the absence and presence of CP [n=4 and p-value is 0.0019 (**)]. (H) Bar graph depicting the change in fluorescence polarization of FAM-labelled HSE upon its interaction with AZD in the presence and absence of CP. Two different concentrations of CP were used in this experiment, as indicated [n=4 and p-value<0.0001 (***)].

To investigate this interaction further, we performed a competition assay using unlabelled (cold) HSE with a sequence identical to that of the FAM-labelled HSE. Incubation of HSE (10 nM) with AZD (10 μM) caused a significant increase in its fluorescence intensity at 520 nm, compared to that of the vehicle (DMSO)-exposed HSE, as expected. However, when the labelled probe-AZD mixture was incubated with an equimolar concentration of the cold probe (10 nM), the fluorescence intensity diminished. A further decrease in fluorescence intensity was observed when a ten-fold molar excess (100 nM) of the cold probe was present in the reaction **(Fig. 6: F&G).** Thus, a dose-dependent decrease in fluorescence intensity of the labelled HSE-AZD mixture was observed in both the above conditions, suggesting that AZD present in the reaction mixture gets distributed between the binding site(s) on the FAM-labelled and unlabelled HSE. The same cold competition assay was also performed by measuring the FP at 520 nm. FP increased considerably upon incubation of labelled HSE with AZD compared to the DMSO-incubated condition. However, increasing concentrations of the cold probe in the reaction caused a significant decrease of FP in a dose-dependent manner **(Fig. 6H).** Thus, the result obtained agreed well with that of the fluorescence intensity study.

We were curious to study the binding affinity of mutant HSE (mHSE) with AZD in the absence of protein and compare it with the affinity of HSE-AZD interaction (mHSE has the same length as that of HSE with key bases mutated, which severely weakens its interaction with HSF1). Interestingly, AZD bound mHSE with a significantly higher affinity compared to that of the HSE-AZD interaction, with a K_d_ of 0.4948 ± 0.2039 μM **(Fig. 6: C, D &E).**

### AZD stimulates the binding of monomeric HSF1 with mutant HSE

If AZD binds mHSE with a higher affinity compared to HSE, does it enhance the binding affinity of mHSE with monomeric HSF1? In our study, AZD induced the multimerization of monomeric HSF1**(Fig.2: G-H)**, but even trimeric/ multimeric HSF1 is reported to bind mHSE with an extremely low affinity [4]. Thus, stimulation of the monomeric HSF1-mHSE binding by AZD might suggest an AZD-induced structural alteration of the DNA molecule.

To address this question, we performed an FP assay to monitor the binding of monomeric HSF1 with FAM-labelled mHSE in the presence and absence of AZD. In the absence of AZD, the protein-DNA interaction was extremely weak, as expected (K_d_: 1172 ± 10.74 nM). But in the presence of AZD (10 μM), the binding affinity dramatically increased with a K_d_ of 84.78 ± 14.19 nM **(Fig. 7: A-C).** Thus, AZD indeed increases the binding affinity of monomeric HSF1 with mHSE considerably, although this affinity remains significantly (∼10 fold) lower than that of the AZD-induced interaction of monomeric HSF1 with HSE (K_d_: 7.98 ± 1.43 nM) **(Fig. 2B &7D).**

**Figure 7:**
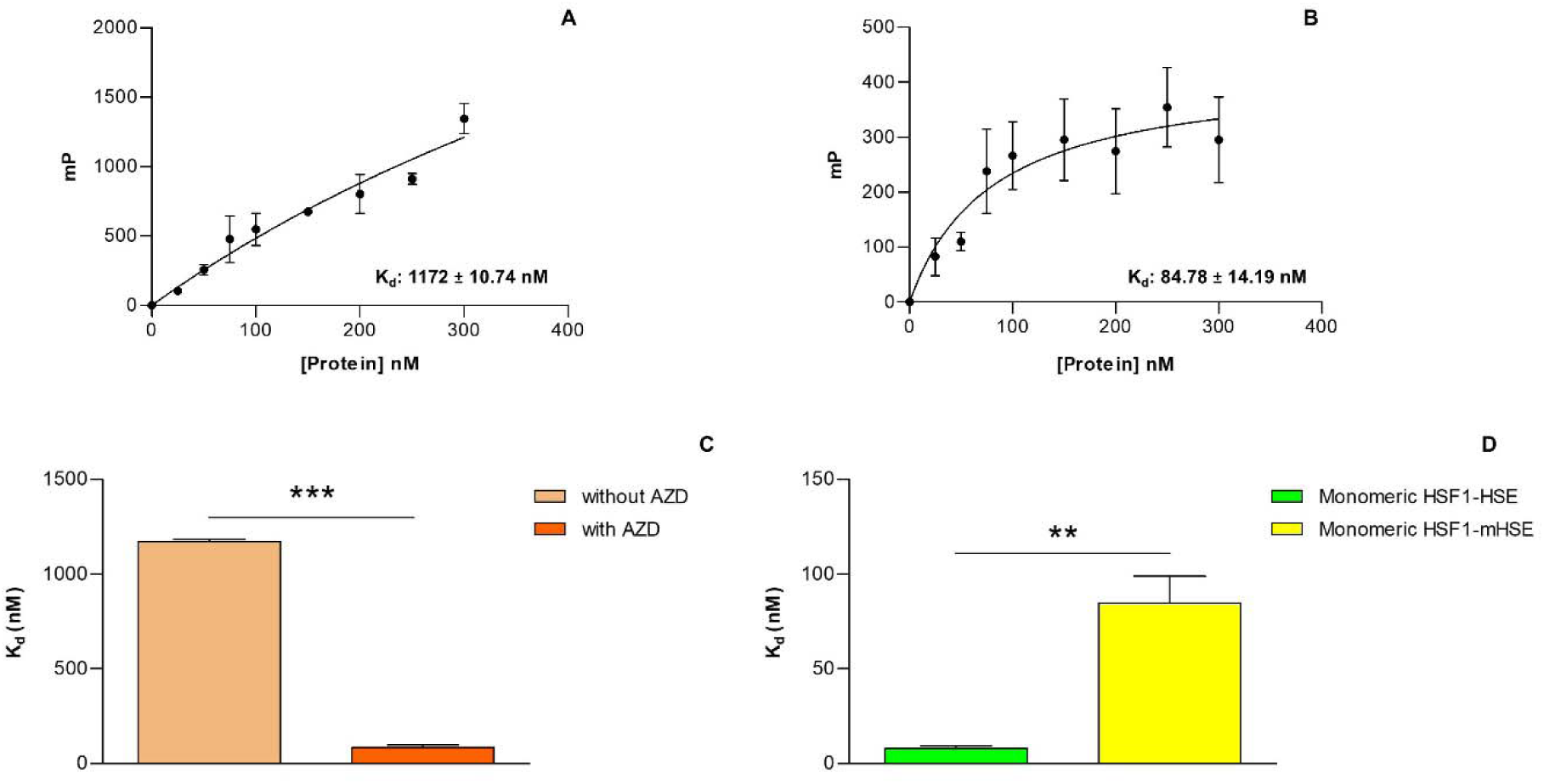
AZD enhances the binding affinity of monomeric HSF1 with mHSE, as demonstrated by FP Assay. Dissociation constants (K_d_) obtained by plotting the millipolarization values of FAM-mHSE against increasing concentrations of monomeric HSF1 in the absence (A) and presence of 10 μM AZD (B). (C) Bar graph comparing the K_d_ for the binding reactions shown in A and B [n=3 and p-value <0.0001 (***)]. (D) Bar graph comparing the K_d_ for the AZD-induced binding of monomeric HSF1 with HSE and mHSE [n=3 and p-value is 0.006 (**)].

### AZD alters the surface hydrophobicity of HSF1 and its derivatives as revealed by 1-Anilinonaphthalene-8-sulfonic acid (ANS) fluorescence analysis

ANS is routinely used as an extrinsic fluorescent probe for detecting exposed hydrophobic patches and cavities on proteins [28]. Protein aggregation is associated with increased exposure of hydrophobic patches. Therefore, an increase in the ANS fluorescence quantum yield under certain treatment conditions in solution suggests a greater ANS incorporation on the surface of proteins. As our DLS results suggested an AZD-induced structural perturbation of the HSF1-WT trimer/ oligomer, HSF1-CT and HSF1-ΔTAD, we tested the alteration in surface hydrophobicity of these proteins upon their incubation with increasing concentrations of AZD, using ANS as the probe.

Exposure of the SEC purified HSF1-WT trimer/ oligomer to increasing concentrations of AZD caused a diminution in the ANS fluorescence quantum yield with a simultaneous shift of the emission maxima to a higher wavelength (red shift). This indicates that the AZD-induced structural destabilization of the HSF1-WT trimer/ oligomer, as indicated by our DLS study, is not caused by the formation of protein aggregates **(Fig. 8: A-C).**

**Figure 8:**
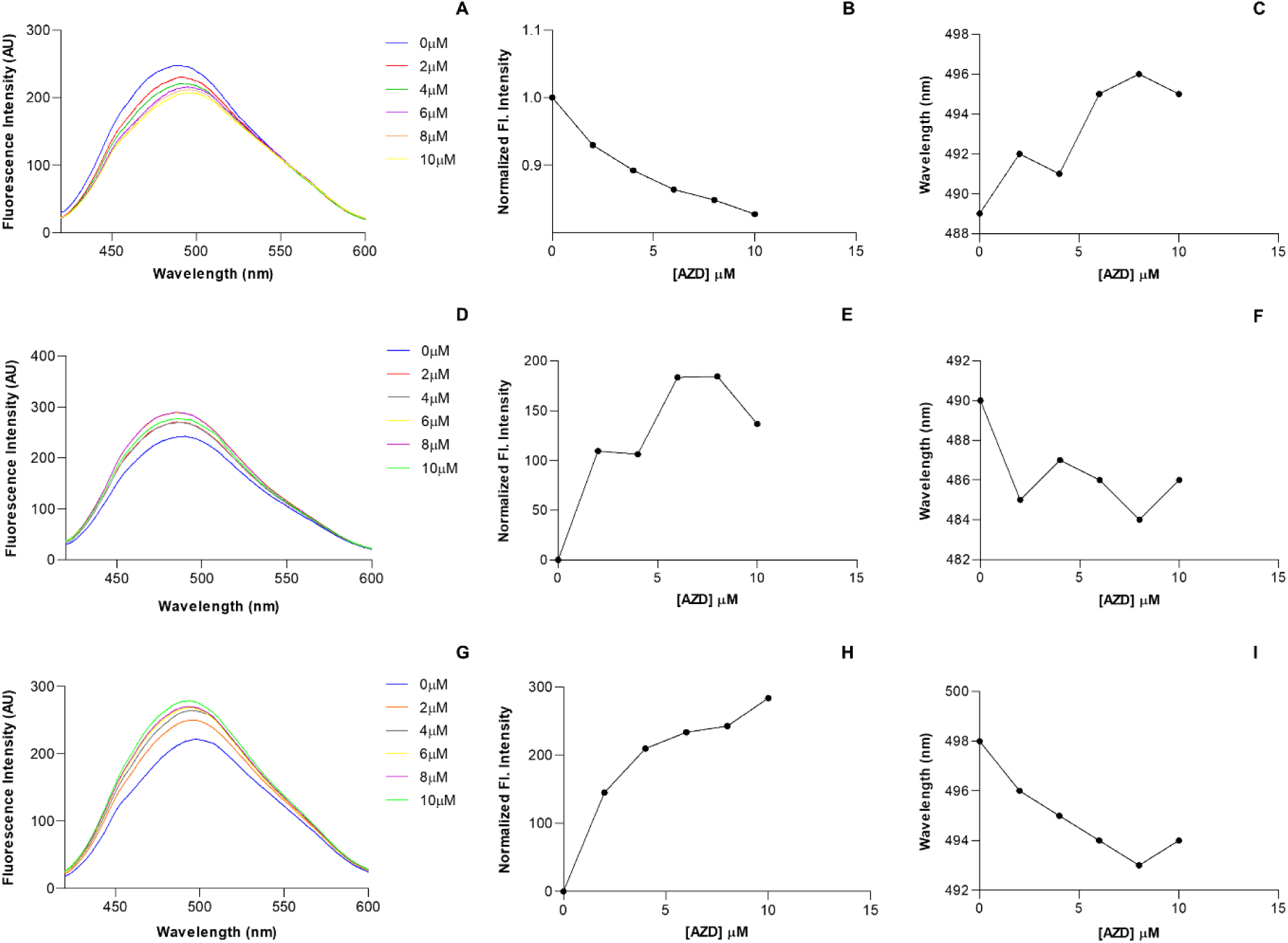
AZD exposure decreases the fluorescence quantum yield of ANS bound to HSF1- WT trimer, while an opposite effect is observed for HSF1-CT and HSF1-ΔTAD. Representative fluorescence emission spectra of ANS bound with HSF1-WT trimer (A), HSF1- CT (D) and HSF1-ΔTAD (G) exposed to increasing concentrations of AZD as indicated. The spectra were obtained at 25°C. Normalized fluorescence intensity (at 490 nm) of ANS bound with HSF1-WT trimer (B), HSF1- CT (E), and HSF1- ΔTAD (H) exposed to increasing concentrations of AZD, as indicated. Change in fluorescence emission maxima (λ_max_) of ANS bound with HSF1-WT trimer (C), HSF1-CT (F), and HSF1-ΔTAD (I) as a function of increasing AZD concentration.

Surprisingly, in contrast to the HSF1-WT trimer/ oligomer, the ANS fluorescence intensity increased considerably when HSF1-CT was exposed to increasing concentrations of AZD, with a concomitant blue shift of the spectral emission maxima **(Fig. 8: D-F).** Similarly, HSF1-ΔTAD, upon incubation with increasing concentrations of AZD, exhibited a significant increase in the ANS fluorescence quantum yield along with a shift of the emission maxima to lower wavelengths **(Fig. 8: G-I).** These results indicate an AZD-induced aggregate formation in these two proteins.

Next, we tested the effect of AZD on the surface hydrophobicity of the monomeric forms of HSF1. Interestingly, incubation of the monomeric form of HSF1-WT with increasing concentrations of AZD revealed an initial increase of the ANS fluorescence quantum yield at lower AZD concentrations, followed by its decrease at higher AZD concentrations. At the highest AZD concentration (10μM), the ANS fluorescence intensity was close to that of the vehicle (DMSO)-exposed protein. The ANS spectral emission maxima exhibited an initial blue shift at lower AZD concentrations, followed by a slight red shift at higher AZD concentrations **(Fig. 9: A-C).** In the case of HSF1-CM, a similar phenomenon was observed, i.e. an initial increase in the ANS fluorescence intensity at lower AZD concentrations was followed by its reduction at higher AZD concentrations. The ANS fluorescence intensity recorded for the AZD (10μM)-exposed protein was close to that of the vehicle-exposed protein. The ANS spectral emission maxima, in this case, exhibited a blue shift at lower AZD concentrations, followed by a redshift at higher AZD concentrations **(Fig. 9: D-F).** Exposure of HSF1-DBD to increasing concentrations of AZD caused an initial rise in the ANS fluorescence intensity, which was followed by its decline at higher AZD concentrations. Incidentally, the ANS quantum yield at the highest AZD concentration (10 μM) was lower than that of the control. For HSF1-DBD, changes in the ANS spectral emission maxima with the increase in AZD concentration followed a pattern like the other two monomeric forms of HSF1 **(Fig. 9: G-I).**

**Figure 9:**
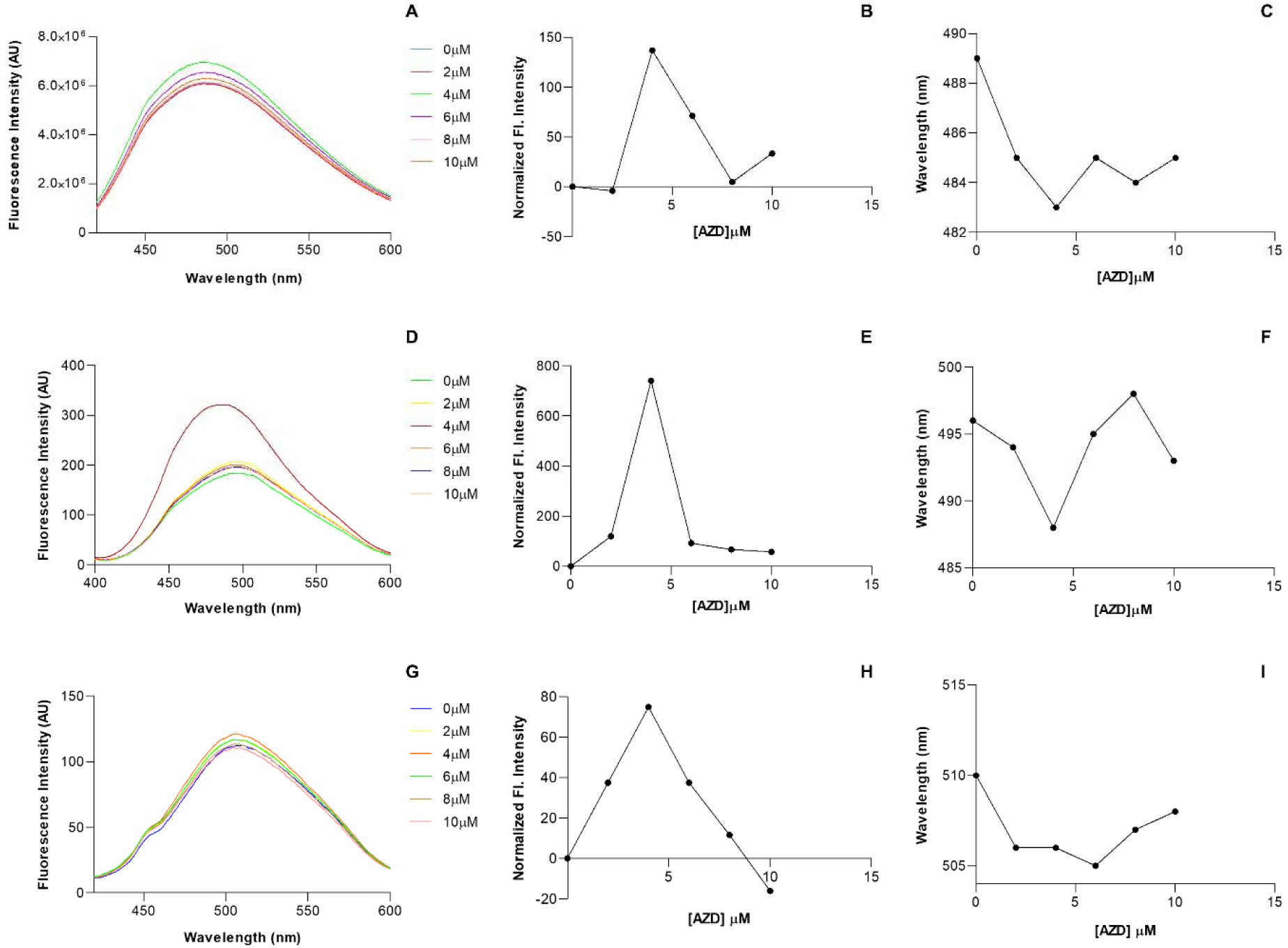
AZD exposure results in a distinct change in the fluorescence quantum yield of ANS bound to HSF1-WT monomer, HSF1-CM, and HSF1-DBD. Representative fluorescence emission spectra of ANS bound with HSF1-WT monomer (A), HSF1-CM (D), and HSF1-DBD (G) exposed to increasing concentrations of AZD, as indicated. The spectra were obtained at 25°C. Normalized fluorescence intensity of ANS bound with HSF1-WT monomer at 485 nm (B), HSF1-CM at 492 nm (E), and HSF1-DBD at 508 nm (H) exposed to increasing concentrations of AZD, as indicated. Change in fluorescence emission maxima (λ_max_) of ANS bound with HSF1-WT monomer (C), HSF1-CM (F), and HSF1-DBD (I) as a function of increasing AZD concentration.

### Analysis of AZD-induced structural perturbation of HSF1-CT and HSF1-**Δ**TAD by Thioflavin T (ThT) fluorescence assay

Our DLS studies and ANS fluorescence assays have indicated the formation of protein aggregates upon treatment of HSF1-CT and HSF1-ΔTAD with AZD. We were interested in investigating the nature of these aggregates. For this purpose, we studied the interaction of ThT, an extrinsic fluorescent dye, with these proteins in the absence and presence of AZD. ThT exhibits a substantial enhancement in its fluorescence intensity upon incubation with characteristic cross-β sheet structures and is thus a suitable probe to characterize amyloid aggregates [33]. When the proteins mentioned above were exposed to increasing concentrations of AZD and subsequently incubated with ThT, a considerable increase in the ThT fluorescence intensity (at 485 nm) was observed **(Fig. 10: A-D).** This indicates that exposure to AZD induced the formation of amyloid-like aggregates of HSF1-CT and HSF1-ΔTAD in a concentration-dependent manner.

**Figure 10:**
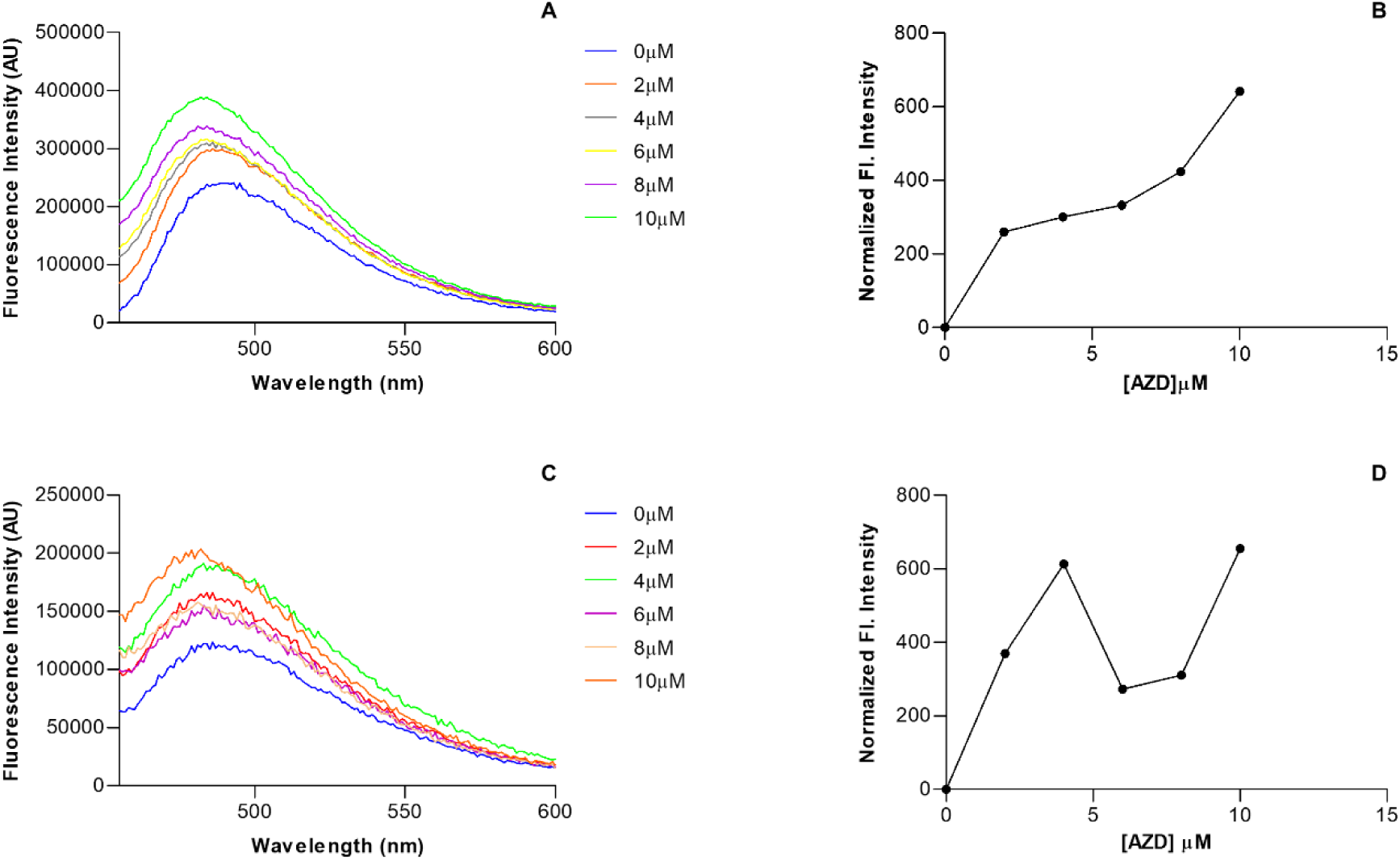
AZD increases cross-β sheet structure of HSF1-CT and HSF1-ΔTAD, as probed by ThT fluorescence assay. Representative fluorescence emission spectra of ThT upon incubation with HSF1-CT (2 μM) exposed to various AZD concentrations, as indicated (A). Normalized fluorescence intensity (at λ_max_= 485 nm) was plotted against AZD concentration for the same experiment in B. Representative fluorescence emission spectra of ThT upon incubation with HSF1-ΔTAD (2 μM) exposed to various AZD concentrations, as indicated (C). Normalized fluorescence intensity (at λ_max_= 485 nm) was plotted against AZD concentration for the same experiment in D.

## Discussion

AZD was previously shown to interact directly with affinity-purified recombinant human HSF1, which increased the protein’s ability to bind HSE. Also, an extract of AZD-treated cells bound HSE with a higher affinity in a gel-shift assay compared to the control sample. These findings were in good agreement with the neuroprotective activities of AZD in *Drosophila* and mouse models of neurodegenerative diseases. Of note, AZD-mediated HSF1 activation is independent of cellular ROS production, as it is known to function as an antioxidant [12, 20, 34].

In the present study, AZD was shown by FP assay to enhance the HSE-binding affinity of the SEC purified monomeric form of HSF1-WT appreciably under non-stress conditions. This was shown to be mediated by the AZD-induced monomer to trimer/oligomer transition of the protein in the complete absence of HSE, as revealed by our DLS analysis **(Fig. 2: A-C, G).** Although HSE binding has been suggested to play some role in inducing trimerization of HSF1-WT monomer *in vitro* [4], in our study, HSE binding did not seem to assist the AZD-induced oligomerization of the HSF1-WT monomer; the fully formed trimer/oligomer bound HSE with high affinity. HSF1 is reported to remain inactive and monomeric in the cytosol. Upon stress exposure, it oligomerizes and moves to the nucleus to engage with its target gene promoters [8]. The results of our *in vitro* studies are consistent with the spatiotemporal separation of HSF1 oligomerization and HSE binding in the cells.

The exact molecular species formed by monomeric HSF1 upon AZD exposure is presently unclear. However, as the multimeric species binds a three-site HSE with high affinity, it seems likely that AZD predominantly triggers the formation of HSF1 trimers. However, the possibility of forming higher order oligomers cannot be ruled out altogether.

Most intriguingly, AZD increased the binding affinity of the constitutively monomeric (CM) form of HSF1 (lacking the oligomerization/ LZ1-3 domain) for HSE by ∼75-fold **(Figs. 1A and 2 D-F).** AZD could induce the oligomerization of this protein independent of HSE; the oligomerized protein bound HSE with an affinity comparable to that of the oligomerized wild-type HSF1 (**Fig. 2: B, E & H)**. Considering the current stress-induced oligomerization domain-dependent activation mechanism of HSF1, this result is completely unanticipated. The wild-type trimeric/ oligomeric HSF1 is known to be held together by inter-subunit hydrophobic contacts and electrostatic interactions in the LZ1-3 domain. As HSF1-CM is devoid of this domain, this naturally raises questions on the point(s) of contact of the individual subunits in the multimeric structure that holds the complex together.

The DBD of HSF1 is its only structurally characterized domain**(Fig. 1A)** [32,35]. The HSF1- DBD exists predominantly in monomeric form in the absence of HSE, as suggested by a low hydrodynamic radius (R_h_) in our DLS studies **(Fig. 4L).** However, a dramatic monomer-to-oligomer shift occurred upon incubation of the protein with AZD, suggested by a substantial increase in R_h_ **(Fig. 4L).** Interestingly, HSF1-DBD was shown to bind HSE in the presence of AZD with a threefold higher affinity compared to its absence. AZD seems to hold the individual DBD subunits in an oligomeric complex, which binds HSE with a higher affinity **(Fig. 4: G-I).** The binding affinity, however, is low compared to the binding affinities of the AZD-induced multimeric forms of HSF1-WT monomer and HSF1-CM. Thus, while the oligomerization domain seems dispensable for AZD-induced multimerization, the interaction of AZD with DBD possibly induces conformational changes in other domains of the HSF1 monomer, enhancing its binding affinity for the HSE.

Our studies on the binding affinities of HSF1-WT monomer, HSF1-CM, and HSF1-DBD with AZD (in the absence of HSE) by protein intrinsic fluorescence spectroscopy revealed very close Kd values **(Fig. 5: A-F).** These results indicate that the DBD is the predominant AZD-binding site in HSF1.

We have tried to interpret our findings in light of the ‘dimer activation model’ proposed by Hentze *et al.* (2016), which explains the stress-mediated HSF1 activation. The LZ4 and LZ1-3 domains interact with each other to stabilize HSF1 in its monomeric form. The larger size of the LZ1-3 domain (75-amino acid residues) compared to the LZ4 domain (42-amino acid residues) allows monomeric HSF1 to form transient dimers via intermolecular interaction with the free region of LZ1-3. The intramolecular interaction tends to destabilize this intermolecular interaction, resulting in the rapid dissociation of such dimers and a predominance of the monomeric form. Interestingly, the intermolecular LZ1-3-LZ1-3 interaction can also destabilize the intramolecular LZ1-3-LZ4 interactions. Thus, anything stabilizing the transient dimer would destabilize the intramolecular LZ1-3-LZ4 interactions. At low HSF1 concentrations, an elevation of temperature leads to the unfolding of LZ4 (and probably of LZ1-3 as well), leading to their detachment from each other. This would promote stabilization of the transient dimer and subsequent HSF1 trimerization [6]. As our findings indicate HSF1-DBD to be the major binding site of AZD in the wild-type HSF1 **(Fig. 5: A-F)** and demonstrate an AZD-triggered DBD oligomerization **(Fig. 4L)**, it seems reasonable that AZD stabilizes the ‘transient dimer’ by interacting with both the DBDs in the dimeric structure, serving as a link between the individual monomeric units. According to the ‘dimer activation model,’ this stabilization could favour the destabilization of the intramolecular LZ1-3-LZ4 interaction of the individual monomers and give enough opportunity for the association of a third monomer with the AZD-stabilized transient dimer at low HSF1 concentrations and low temperatures. This results in the displacement of LZ4 from LZ1-3, forming a stable trimeric structure via the LZ1-3 domains **(Fig. 11).**

**Figure 11:**
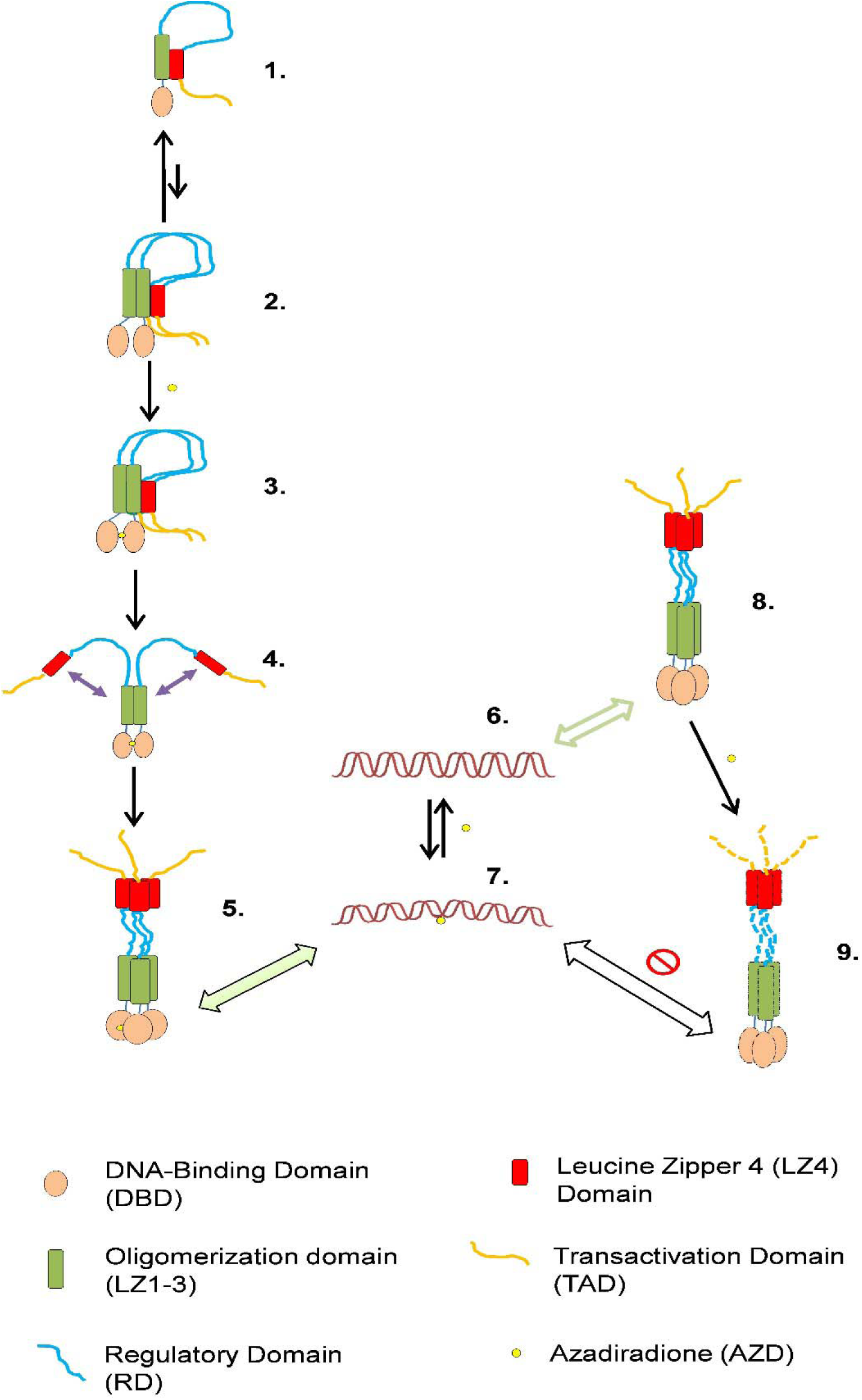
A plausible model for the context-dependent AZD-mediated activation and inhibition of human HSF1. (1) Monomeric HSF1, where LZ1-3 and LZ4 domains remain associated. **(2)** A transient dimer, formed by the association of two such monomers, quickly reverts to the monomeric form. **(3)** AZD stabilizes the transient dimer, via contact with the DBDs of the individual monomers. **(4)** Transient dimer stabilisation leads to destabilisation of the LZ4-LZ1-3 interactions of the individual monomers, shown by purple arrows. **(5)** Association of a 3^rd^ monomer with the dimeric structure forms the active trimeric HSF1, with AZD bound. **(6)** HSE. **(7)** AZD exposure induces conformational alteration(s) (possible bending) of HSE. This AZD-bound HSE binds AZD-trimerized HSF1 (5) with a very high affinity, as represented by the green arrow. **(8)** Pre-assembled trimeric/ oligomeric form of HSF1, which binds HSE (6) with high affinity in the absence of AZD. **(9)** The presence of AZD causes structural destabilisation/ aggregation of the pre-assembled trimeric/ oligomeric HSF1, reducing its affinity for HSE, as represented by the white arrow.

Strikingly, our studies show that AZD triggers the oligomerization and HSE-binding of HSF1- ΔLZ1-3 with an efficiency comparable to that of HSF1-WT monomer **(Fig. 2).** Here, AZD-mediated inter-subunit DBD-DBD interactions seem to hold the multimer together. A winged helix-turn-helix fold was identified in the HSF-DBDs from *Kluyveromyces lactis*, *Drosophila*, and human [36–39]. Unlike many other transcription factors possessing a wing motif, the wing motif of HSF1-DBD does not contact the DNA. Rather, it mediates protein-protein interactions within the same trimeric unit; the HSE-bound DBDs interact via this wing motif. The wing motif also mediates interactions between the HSE-bound DBD molecules from adjacent trimeric units, thereby facilitating a synergistic binding of DBDs to HSEs having multiple binding sites [35]. Our results are consistent with the idea that AZD facilitates interactions among the DBDs of HSF1 subunits through this winged helix-turn-helix fold either directly or indirectly. Future studies will test this hypothesis.

Surprisingly, we observed an inhibitory effect of AZD on the pre-assembled oligomeric forms of HSF1. Both the WT trimer/ oligomer (obtained by SEC of affinity-purified HSF1-WT) and the constitutively trimeric/ oligomeric mutant form (HSF1-CT) bound the HSE with lowered affinity in the presence of AZD in comparison to its absence **(Fig. 3: A-F).** DLS analyses of these proteins in the absence of HSE revealed an AZD-induced slight increase in their hydrodynamic radii. This could be attributed to a partial destabilization of the proteins’ structure upon incubation with AZD, which could explain the observed decrease in HSE-binding affinities **(Fig. 3: G & H).** Our results suggest that the pre-assembled HSF1 trimer/ oligomer present in the cytoplasm/ nucleus is functionally inhibited by AZD, in contrast to the AZD-mediated functional activation of monomeric HSF1.

Cancer cells rely on HSF1 for their survival and proliferation and hence maintain the protein in a constitutively active state [14,15]. Therefore, inhibition of oligomeric HSF1 in cancer cells could sensitize these cells to standard anticancer drugs. As such, HSF1 inhibition has become a promising strategy in anticancer drug discovery, although transcription factors are traditionally considered “undruggable” [40]. HSF1-targeting drugs are not expected to affect the survival of normal cells, as HSF1-target genes are supposed to be non-essential for their growth and survival under stress-free conditions. Indeed, *hsf1-/-* mice are viable under controlled laboratory conditions and display a lower incidence of tumours[14].

Several HSF1 inhibitors were reported to directly interact with HSF1. DTHIB, a synthetic compound, interacts with HSF1 directly and facilitates selective degradation of nuclear HSF1 without affecting its HSE-binding ability in *vitro*[41]. I_HSF1_115 was reported to block HSF1’s transcriptional activity by interfering with ATF1-containing complex assembly[42]. Peptide 14, which mimics a discrete region of the LZ4 domain, interacts with the LZ1-3 domain to suppress the DNA binding of HSF1 without altering its multimeric structure [43]. The inhibitory effect of AZD, however, seems to be directed only at the pre-assembled homomultimeric form of HSF1, which is an intermediate in its cellular activation-deactivation pathway. Thus, AZD acts both as an activator and inhibitor of HSF1 in a context-dependent manner, revealing its ameliorating effects for both neurodegenerative diseases [12] and cancers. Indeed, El-Senduny *et al.* (2021) reported the anticancer activity of AZD on MDA-MB-231, the triple-negative breast cancer cell line. AZD-driven HSF1 inhibition, however, was not explored in that study [44].

How does the same compound enhance the activity of the monomeric forms of HSF1 and hinder the activity of its multimeric forms? As per our DLS analyses, AZD does not dissociate the pre-assembled oligomeric forms of full-length HSF1 to individual monomers **(Fig. 3: G & H).** To further explore the cause behind AZD-induced inhibition of oligomeric HSF1, we studied the binding of the fluorescent probe ANS with these proteins treated with increasing concentrations of AZD. The treatment of HSF1-CT with increasing AZD concentrations caused a rise in the ANS fluorescence intensity signal with a concomitant blue shift of the fluorescence emission maxima, suggesting an AZD-induced increase in the exposure of hydrophobic patches on the protein’s surface, leading to aggregation. This protein aggregate was shown to be amyloid-like by ThT fluorescence assay. **(Fig. 8: D-F**; **Fig. 10: A & B).** HSF1-ΔTAD, essentially multimeric, also exhibited similar behaviour as HSF1-CT, demonstrating that deletion of the TAD renders HSF1 aggregation-prone **(Fig. 8: G-I**; **Fig. 10: C & D).** Notably, contrary to our present observations, AZD was previously reported to completely destabilize Tau aggregates and inhibit the aggregation of Tau protein [21]. In the case of the HSF1-WT trimer/ oligomer, surprisingly, a decrease in the ANS fluorescence intensity was observed upon incubation of the protein with increasing concentrations of AZD, accompanied by a red shift of the spectral emission maxima. It should be remembered that both hydrophobic and electrostatic interactions have been implicated in the binding of ANS with proteins: the negatively charged sulfonate group interacting with positively charged amino acid residues and the aromatic rings with hydrophobic residues in an oriented manner [33]. Thus, it is reasonable to infer that the destabilization of HSF1-WT trimer/ oligomer induced by AZD causes a loss of complementarity of the positively charged and hydrophobic residues, which is essential for the binding of ANS. As a result, ANS binds the protein loosely via only one form of interaction and hence gets detached easily, accounting for the decrease in fluorescence quantum yield and the spectral redshift **(Fig. 8: A-C).**

Sengupta *et al.* (2023) correlated the amyloid formation of the mutant and wild-type forms of p53 in cancer tissues with the loss of DNA binding and transcriptional activities of the transcription factor and increased cancer grades. Misfolding and/ or aggregation and subsequent amyloid formation of p53 compromise its function as a tumour suppressor [45]. It is likely, therefore, that AZD-induced amyloid formation in HSF1-CT and HSF1-ΔTAD, as demonstrated in our study, is responsible for the observed decrease in their HSE-binding affinities. AZD-induced amyloid formation of HSF1 in cancer cells and the consequent loss of its DNA-binding activity could be one of the reasons behind the demonstrated anticancer properties of AZD [44]. Studies on the interaction of ANS with HSF1-WT monomer and HSF1-CM treated with increasing concentrations of AZD revealed that a gradual increase in the concentration of AZD led to an initial rise in the ANS fluorescence quantum yield followed by a decline of the same **(Fig. 9: A-F).** This was accompanied by a blue shift of the ANS spectral emission maxima at lower concentrations of AZD, followed by its redshift at higher concentrations. This indicates an initial destabilization of the proteins’ structure (possible aggregation) at lower AZD concentrations followed by stabilization at higher concentrations. HSF1-DBD also showed a similar behaviour **(Fig. 9: G-I).** Structural stabilization of the proteins at high AZD concentration is corroborated by the fact that 10 μM AZD (the highest AZD concentration used in the ANS binding assays) has been shown to increase their DNA binding affinities appreciably. A similar observation was reported in a previous study, where maximal aggregation of conalbumin occurred at lower percentages of fluoroalcohols, while higher percentages caused an increase in the protein’s helical propensity [46].

Further, our studies revealed the specific and reversible interaction of AZD with HSE *in vitro* in the absence of protein **(Fig. 6: A, B, F, G & H).** Does this interaction play a role in the AZD-induced HSE binding by monomeric HSF1? To address this question, first, we studied the binding of AZD with mutant HSE (mHSE) under the same conditions as that of HSE. Of note, both the monomeric and oligomeric forms of HSF1 were reported to bind mHSE with extremely low affinities, owing to the mutation of key bases involved in HSF1 binding [4]. Interestingly, AZD was found to bind mHSE with a ∼2.6-fold higher affinity than HSE under the same reaction conditions **(Fig. 6: C-E).** Next, to investigate the possible role of mHSE-AZD interaction in the binding of mHSE with monomeric HSF1, the same FP assay described previously for HSF1-HSE binding was performed. The affinity of monomeric HSF1-mHSE interaction was raised ∼14-fold in the presence of AZD compared to its absence **(Fig. 7: A-C).** The binding affinity still, however, remained appreciably lower than that of monomeric HSF1- HSE interaction in the presence of AZD **(Fig. 7D).** This observation suggests that, apart from inducing multimerization of monomeric HSF1 (as shown by DLS assay, **Fig. 2G**), AZD induces conformational alteration(s) in the DNA molecules to add to the binding affinities. The lowered binding affinity of multimerized HSF1 with mHSE compared to that of multimerized HSF1 with HSE reflects the effect of mutation of critical bases that are vital for HSF1-HSE interaction [4]. This AZD-driven conformational alteration(s) of DNA could explain the stronger binding of AZD-multimerized HSF1 with HSE (K_d_: 7.98 ± 1.43 nM), compared to the binding of pre-assembled HSF1-WT trimer and HSF1-CT with HSE (K_d_: 17.13 ± 6.278 nM and 25.07 ± 8.094 nM respectively) **(Figs. 2B, 3A & 3D).**

The crystal structure of the HSE-bound HSF1 trimer consisting of three DBDs suggests the preferential recognition and binding of a curved DNA molecule by HSF1[35]. It thus seems possible that AZD induces curvature in both the HSE and mHSE oligonucleotides, contributing to the enhanced binding affinity of AZD-multimerized HSF1 with these DNA molecules.

## Conclusion

Our studies revealed that AZD could influence the HSE-binding activity of HSF1 depending on the molecular state of the protein. While AZD efficiently enhanced the HSE-binding activity of purified monomeric HSF1 through oligomerization, it apparently hindered the HSE-binding of the pre-assembled oligomeric forms of HSF1. Unlike monomeric HSF1, the pre-assembled constitutively trimeric/ oligomeric HSF1 transitioned to amyloid-like aggregates in the presence of AZD. AZD drove the oligomerization of the HSF1 monomer by interacting with its DNA-binding domain without involving the oligomerization domain of the protein, which is known to be indispensable for stress-induced HSF1 activation. Furthermore, AZD-induced conformational changes in the HSE facilitated monomeric HSF1 binding. Taken together, our studies have revealed a unique mechanism of context-dependent HSF1 activation and inhibition by AZD, a triterpenoid that is non-toxic to healthy cells.

## Materials and methods

### Materials and reagents

All organic solvents and silica gel (230-400 mesh) used for AZD purification and characterization, molecular weight standards for size-exclusion chromatography (Sigma), DNA oligonucleotides (HPLC grade), sodium chloride (NaCl), imidazole, tricine, 6-aminohexanoic acid, sodium dodecyl sulfate (SDS), glycerol, dithiothreitol (DTT), and dimethyl sulfoxide (DMSO) were purchased from Sigma-Aldrich (St. Louis, MO). Luria broth, ampicillin, and isopropyl-β-D-thiogalactopyranoside (IPTG) were purchased from HiMedia Laboratories, LLC. 4-(2-hydroxyethyl) piperazine-1-ethanesulfonic acid (HEPES), phenylmethylsulfonyl fluoride (PMSF), lysozyme (3X crystallized), tris base, glycine N, N, N′, N′-tetramethyl ethylenediamine (TEMED), acrylamide, bisacrylamide, ammonium persulfate (APS), Coomassie brilliant blue G250 and R250, 8-anilinonaphthalene-1-sulfonic acid (ANS), and thioflavin T (ThT) were purchased from Sisco Research Laboratories (SRL) Pvt. Ltd. Nickel-nitrilotriacetic acid (Ni-NTA) agarose beads were purchased from Qiagen (Germany). Bradford reagent was purchased from Bio-Rad. Restriction enzymes, T4 DNA ligase, and Q5 DNA polymerase were purchased from New England Biolabs (NEB). A pre-stained protein ladder was purchased from Abcam. Dialysis membrane (12-14 kDa MW cutoff) was purchased from SpectraPor®.

### Purification and characterization of Azadiradione

Azadiradione (AZD) (approx. 800 mg) was purified from 5 kg neem seed powder, following the method described by Nelson *et al.*[20]. Thin-layer chromatography (TLC) and analytical high-performance liquid chromatography (HPLC) were used to estimate the purity of the compound, which was found to be ∼95 % **(Fig. S1: A & B).** The identity of the compound was further confirmed by Nuclear Magnetic Resonance (NMR) spectroscopy (^1^H-NMR, ^13^C-NMR, and DEPT135-NMR) **[Fig. S1: C, D & E].**

### Expression and purification of His_6_-tagged human HSF1 and its derivatives

To investigate the interaction of AZD with human HSF1, the following HSF1 derivatives were used **(Fig. 1A).** In HSF1-ΔLZ1-3, the oligomerization domain of HSF1 has been deleted, consequently, this protein remains constitutively monomeric (CM)[4]. Three residues in the LZ4 domain have been mutated in HSF1-LZ4mutant (LZ4m) to disrupt its interaction with the LZ1-3 or oligomerization domain (M391K, L395P, and L398R). This protein, therefore, remains constitutively trimeric/ oligomeric (CT) [4,22,23]. The HSF1-DNA Binding Domain or HSF1- DBD, consists of residues 1-123 [4]. HSF1-ΔLZ4-TAD consists of the DNA-binding, oligomerization, and regulatory domains, and HSF1-ΔTAD consists of the DNA-binding, oligomerization, regulatory, and LZ4 domains.

We also made several attempts to purify a truncation derivative of HSF1 consisting of the DBD and oligomerization domain (amino acid residues 1-203), but this derivative precipitated out of solution due to aggregation during dialysis, presumably owing to the high level of hydrophobicity in the solution-exposed regions. Hence, it could not be used in the present study. The codon-optimized DNA fragments encoding wild-type human HSF1 (HSF1-WT) and its mutants HSF1-ΔLZ1-3 and HSF1-LZ4m cloned in pET15b expression vector at *NdeI/XhoI* restriction sites were kind gifts from Prof. Dennis J. Thiele (Sisu Pharma). The C-terminal truncation derivatives: HSF1-DBD (amino acid residues 1-123), HSF1-ΔLZ4-TAD (amino acid residues 1-383), and HSF1-ΔTAD (amino acid residues 1-409) were constructed similarly in the pET15b vector, their coding sequences being PCR-amplified from the codon-optimized *hsf1* sequence. These constructs were transformed into the *E. coli* strain BL21 (DE3) to overexpress the proteins in secondary cultures (OD_600_∼0.5) using 0.5 mM IPTG for 2-3 h at 28°C.

The cell pellets obtained by centrifugation of the IPTG-induced cultures were resuspended in an ice-cold Lysis Buffer [50 mM HEPES-NaOH (pH 7.4), 300 mM NaCl, 25 mM imidazole (pH 8), 10% (v/v) glycerol,1 mM PMSF and 0.4 mg/ml lysozyme] for 30 min followed by sonication for 5 min in an ice-bath sonicator (30-sec bursts with 60-sec gaps). The crude lysate thus obtained was immediately centrifuged at 12000 rpm for 30 min at 4°C to collect the supernatant (cleared lysate) in a fresh tube. As HSF1 is highly sensitive to proteases, the following steps were performed in a cold room using ice-cold buffers. The cleared lysate was mixed with 2 ml of Ni-NTA agarose beads (pre-equilibrated in Lysis Buffer) per liter of secondary bacterial culture and rotated in a Rotospin for 2 h. This suspension was transferred to a column, and the flowthrough was collected under gravity. The settled beads in the column were washed once with a 5× bead volume of Wash Buffer [50 mM HEPES-NaOH (pH 7.4), 300 mM NaCl, 30 mM imidazole (pH 8), 10% (v/v) glycerol], followed by the elution of bead-bound proteins with an Elution Buffer [50 mM HEPES-NaOH (pH 7.4), 300 mM NaCl, 250 mM imidazole (pH 8), 10% (v/v) glycerol]. The eluted proteins were collected in ∼200 μl fractions. The purity and integrity of the protein-containing fractions were analyzed by reducing SDS-PAGE followed by Coomassie Brilliant Blue R250 gel staining. The SDS-PAGE profile of all the proteins showed a single major band with minimal degradation **(Fig. S2: A-G).** The fractions containing higher concentrations of the desired protein were pooled together and dialyzed overnight against Dialysis Buffer [25 mM HEPES-NaOH (pH 7.4), 150 mM NaCl, and 10% (v/v) glycerol].

### Size Exclusion Chromatography of Affinity Purified Proteins

The oligomeric status of the affinity-purified proteins was determined by Size Exclusion Chromatography (SEC) using a Superdex200 Increase 10/300 GL column (Cytiva) pre-calibrated with standard molecular weight markers for SEC run in an KTA pure^TM^ chromatography system. A standard curve was obtained by plotting these markers’ log [Molecular weight] versus Elution Volumes. The column was run at a flow rate of 0.5 ml/min using Dialysis Buffer as the mobile phase. The elution profile of each protein preparation was monitored by associated absorbance at 280 nm. The eluted proteins were collected in 200 μl fractions using a fraction collector. The protein concentration of each fraction was estimated by Bradford’s method. Subsequently, aliquots from these fractions were analysed by Blue Native PAGE. The fractions were finally flash-frozen in liquid nitrogen and stored at -80°C for further studies [6].

### Blue Native PAGE

Blue Native PAGE was performed with the monomeric and multimeric molecular species separated by SEC following the protocol **Wittig *et al.* (2006)** described, with some modifications[24]. Briefly, protein samples were mixed with 10X Sample Buffer [glycerol 50% (v/v), Coomassie Brilliant Blue G250 solution 0.2%, and Cathode Buffer-B 10% (v/v) **\***] and resolved in a 7% polyacrylamide native gel using Cathode Buffer-B [50 mM Tricine, 7.5 mM Imidazole (pH 7)] in the upper tank and Anode Buffer [25 mM Imidazole (pH 7)] in the lower tank at 4°C for 2-4 h. The gel was run at a constant current (15 mA) except at 100 V in the beginning for allowing the protein samples to enter the gel. It was monitored that the voltage did not exceed 500V for a gel dimension of 0.16 × 14 × 14 cm that was used. Protein bands were readily visible in the gel due to the Coomassie Brilliant Blue G250 dye in the sample buffer. [**\***Cathode Buffer-B 10% (v/v) means 1 ml Cathode Buffer-B in a total volume of 10 ml Sample Buffer (10X)]

### Fluorescence Polarization (FP) Assay

To study the interaction of human HSF1 and its derivatives with the canonical three-site heat shock element (hereafter referred to as ‘HSE’), the DNA sequence 5’- CCTG<u>GAA</u>TA<u>TTC</u>CC<u>GAA</u>CTGGC-3’ containing three inverted 5’-nGAAn-3’ repeats tagged with the fluorophore fluorescein amide (FAM) at its 5’ end was used [4]. To obtain a double-stranded DNA probe, the tagged sequence and its untagged complementary sequence were annealed by heating them together at equimolar concentrations in an annealing buffer (5X annealing buffer composition: 50 mM Tris pH 8, 0.5 mM EDTA, 0.75 M NaCl) at 95°C in a heat block for 5 min, followed by slow cooling to room temperature.

1 nM of the annealed probe was titrated with increasing concentrations of HSF1 and its derivatives in the absence and presence of AZD (10 μM). Fluorescence polarization was measured at 25°C using an excitation wavelength (λ_ex_) of 495 nm and an emission wavelength (λ_em_) of 517 nm. For this, the fluorescence intensities (I) at 517 nm were measured for each sample in the following four orientations of the excitation and emission polarizers: HH, HV, VH, VV (in each case, the first and second letters indicate the relative positions of the excitation and emission polarizer, respectively; H: Horizontal, V: Vertical). Fluorescence Polarization (P) was calculated for each sample using the following equation [25]:

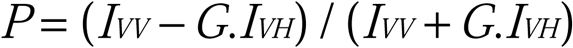

Here, G (Grating correction factor) = *I_HV_* / *I_HH_*

Protein-DNA binding curves for HSF1 and its derivatives were obtained by plotting milli- polarization (mP) values for increasing protein concentrations, and equilibrium dissociation constants (K_d_) were determined for all the protein-DNA interactions using a ‘one site-specific binding’ fit of the curves in GraphPad Prism 5. The interaction of monomeric HSF1 with mutant HSE (mHSE) in the presence and absence of AZD was studied similarly. The mHSE sequence is 5’-CCTG**G<u>CG</u>**TA**<u>G</u>TC**CC**<u>CGC</u>**CTGGC-3’. (The underlined represent the mutated bases) [4].

### Dynamic Light Scattering (DLS) Assay

To compare the oligomeric status of human HSF1 and its derivatives in the presence and absence of AZD, DLS assays were performed using a Zetasizer Nano S (Malvern Instruments, Malvern, UK). The protein samples in the dialysis buffer were passed through a 0.22 μm syringe filter to remove any particulate matter before performing the assays. Samples (protein: ligand molar ratio set as 1:5) were incubated for 10 min at 25°C and scanned for DLS. Each data was taken as an average of 10 scan results. The hydrodynamic radius (R_H_) was calculated by the instrument’s software using the Stokes-Einstein equation:

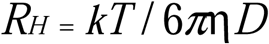

where k is Boltzmann’s constant, T is the absolute temperature, is the medium’s viscosity, and D is the translational diffusion coefficient of the particle under study [26].

### Protein Intrinsic Fluorescence Spectroscopy

The interactions of AZD with HSF1-WT monomer, HSF1-CM, and HSF1-DBD were studied in the absence of HSE, employing the intrinsic fluorescence of these proteins. The studies were conducted in the previously mentioned dialysis buffer. 2 μM of each protein was incubated with increasing concentrations (0-10 μM) of AZD at 4°C. DMSO was kept at 2% (v/v) for each reaction mixture. Next, the samples were excited at 280 nm, and the emission spectra were recorded from 300 to 400 nm at 25°C, keeping the excitation and emission slit widths at 5 nm. The fluorescence spectrum of AZD in the buffer was subtracted from the protein spectra. For each protein, the fluorescence intensity values at the maximum emission wavelength (λ_max_) for each AZD concentration were derived from the spectra and plotted against AZD concentrations to obtain the binding isotherms. Data were analyzed in GraphPad Prism 5 using nonlinear regression with a ‘one site-specific binding’ model to obtain the K_d_ [27].

### Analysis of HSE-AZD interactions by fluorescence spectroscopy

The interactions of AZD with HSE and mHSE were studied in dialysis buffer. 10 nM 5’-FAM labelled double-stranded oligonucleotides mentioned previously were incubated with 0-10 μM of AZD at 4°C, keeping a 2% (v/v) DMSO concentration in all the reaction mixtures. Fluorescence emission spectra of the samples were recorded at 25°C from 505-560 nm upon their excitation at 495 nm, setting the excitation and emission slit widths at 5 nm. For each oligonucleotide, the fluorescence intensity values at 520 nm (λ_max_) were plotted against increasing AZD concentrations to obtain the binding isotherms. The K_d_ were obtained by analyzing the data in GraphPad Prism 5 using nonlinear regression with a ‘one site-specific binding’ model [4,27].

### Cold competition assay for analysing the interaction of HSE with AZD

Four reactions were set up at 4°C in dialysis buffer as follows, maintaining DMSO concentration at 2% (v/v): (a) 10 nM FAM-HSE + DMSO (vehicle), (b) 10 nM FAM-HSE + 10 μM AZD, (c) 10 nM FAM-HSE + 10 nM cold HSE + 10 μM AZD, (d) 10 nM FAM-HSE + 100 nM cold HSE

+ 10 μM AZD. Following incubation, the fluorescence intensities of the samples were recorded at 25°C with emission in the range of 505-560 nm following excitation at 495 nm. The fluorescence intensities at 520 nm for all the samples were plotted in a bar graph using GraphPad Prism 5 and compared. The fluorescence polarization (FP) of FAM-HSE was analyzed under identical conditions, as elaborated above. Here, the FP of the samples at 520 nm were recorded after excitation at 495 nm and plotted and compared in a bar graph.

### 8-Anilinonaphthalene-1-sulfonic acid (ANS) Binding Assay

The proteins (2 μM) were incubated with varying concentrations of AZD (0-10 μM) in dialysis buffer for 30 min at 4°C in the dark with DMSO maintained at 2% (v/v) in each reaction mixture. Next, the ANS stock solution prepared in dialysis buffer was added at a final concentration of 40 μM (20-fold molar excess of proteins) to the reaction mixtures, followed by a further incubation of 30 minutes in the dark at 4°C. After this, the emission spectra of the samples were recorded from 420-600 nm at 25°C following excitation at 390 nm [28], setting both the excitation and emission slit widths at 5 nm. The fluorescence intensity values at the maximum emission wavelength (λ_max_) for each protein were plotted against the AZD concentrations.

### Thioflavin T (ThT) Binding Assay

Proteins (2 μM) were incubated with increasing concentrations of AZD (0-10 μM) in dialysis buffer [maintaining 2% (v/v) DMSO] for 30 minutes in the dark at 4°C. ThT stock solution was prepared in dialysis buffer and added at a final concentration of 4 μM to the reaction mixtures (molar ratio of protein: ThT was 1:2), followed by a further incubation of 30 minutes in the dark at 4°C. After this, the emission spectra of the reactions were recorded from 450-600 nm at 25°C following excitation at 440 nm [29]. Appropriate blanks were prepared by incubating AZD at varying concentrations with 4 μM ThT in the dialysis buffer, and their emission spectra were subtracted from the sample spectra. The fluorescence intensity values at λ_max_ were plotted against AZD concentrations.

### Statistical Analyses

The results are presented as mean ± SEM. Data analyses were done using one-way ANOVA (Tukey’s Multiple Comparison Test, comparing more than two groups) and two-tailed unpaired t-tests (comparing two groups) in GraphPad Prism 5 software. The results were statistically significant at P < 0.05.

## Acknowledgements

The authors thank Prof. Gopal Chakrabarti for reading and critically commenting on the manuscript. Thanks are due to Mr. Dipak Konar, Mr. Swaroop Biswas, Mr. Smriti Ranjan Maji and Mr. Mrinal Das for their assistance during the experiments. AM was the recipient of Junior and Senior Research Fellowships from the Dept. of Biotechnology (DBT), Govt. of India. PB was the recipient of a Research Associateship from Bose Institute. CM was the recipient of DST-Inspire Fellowship, Govt. of India. This work was funded by Bose Institute and DBT.

## Author contributions

AM: performed most of the experiments, analyzed the data & wrote the manuscript; PB: cloned the HSF1 variants, analyzed the data, reviewed & edited the manuscript; CM & AM: purified & characterized AZD; NH: performed one FP assay; JM & SCM: reviewed the manuscript; KJ: resources; MP: conceptualized, designed & supervised the study, acquired funding, reviewed & edited the manuscript.

## Abbreviations

HSF1: Heat Shock Factor 1
HSE: Heat Shock Element
AZD: Azadiradione
DBD: DNA- Binding Domain
LZ: Leucine Zipper
RD: Regulatory Domai
TAD: Transactivation Domain
FAM: Fluorescein Amide
FP: Fluorescence Polarization
DLS: Dynamic Light Scattering
CP: Cold Probe
ANS: 8-Anilinonaphthalene-1-sulfonic acid
ThT: Thioflavin T.

## Conflict of interest

The authors declare no conflict of interest.

## Notes

### Competing Interest Statement

The authors have declared no competing interest.

