## Supplementary information for "Azadiradione regulates Heat Shock Factor 1 function by interacting with its DNA-binding domain independent of the oligomerization domain"

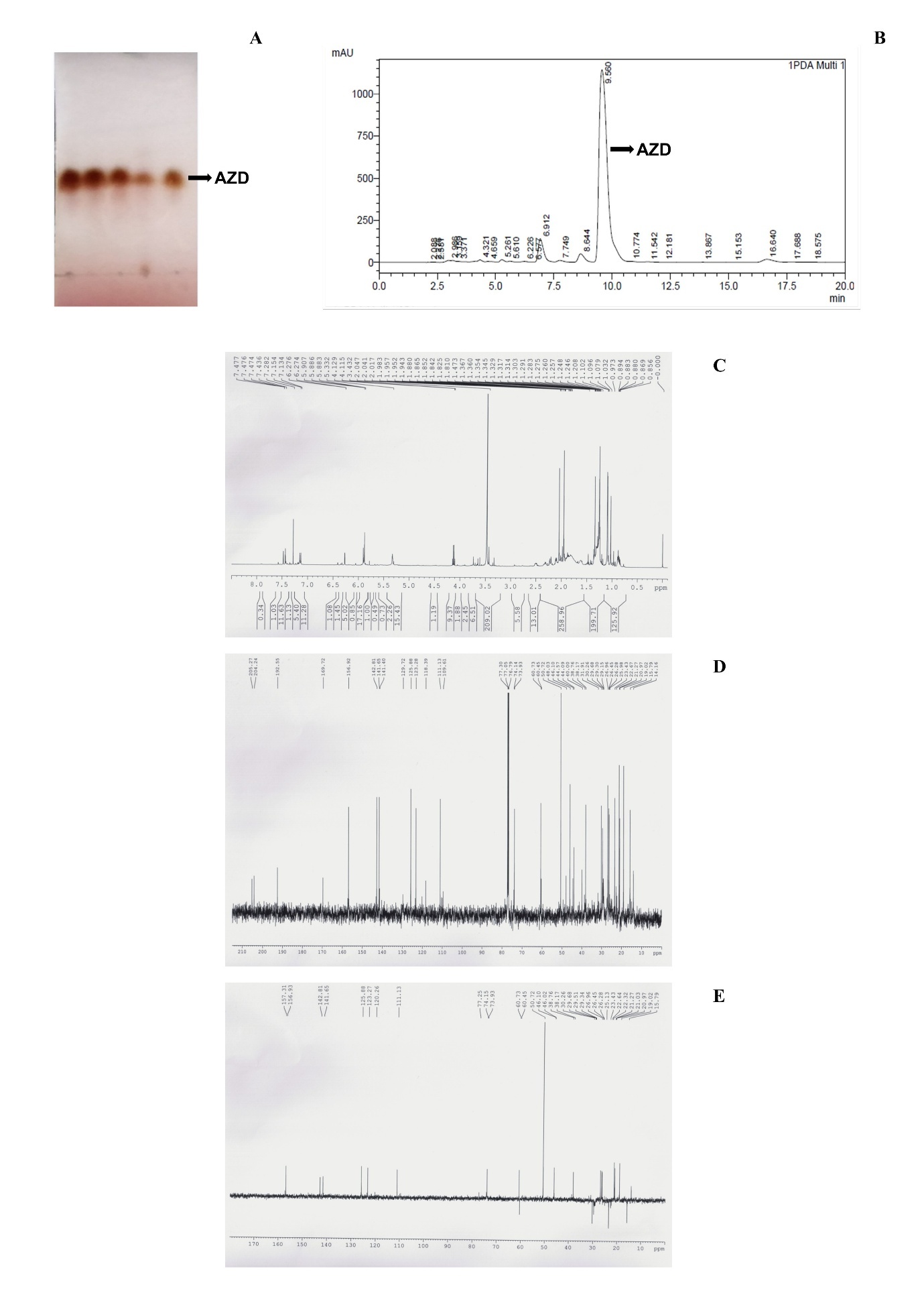

**Fig. S1. Purification and characterization of Azadiradione (AZD).**

Thin Layer Chromatography (TLC) profile of an AZD preparation. TLC was performed using a silica gel coated TLC plate (Merck) with mobile phase Hexane: Ethyl acetate :: 4:1 (A).

Analytical High-Performance Liquid Chromatography (HPLC) profile of an AZD preparation. HPLC was performed using a C-18 column with a mixture of acetonitrile (60%) and water (40%) as the mobile phase. The AZD peak having a retention time of 9.56 min is indicated by an arrow (B).

^1^H-NMR (C), ^13^C-NMR (D) and DEPT-135 NMR (E) spectra of AZD.

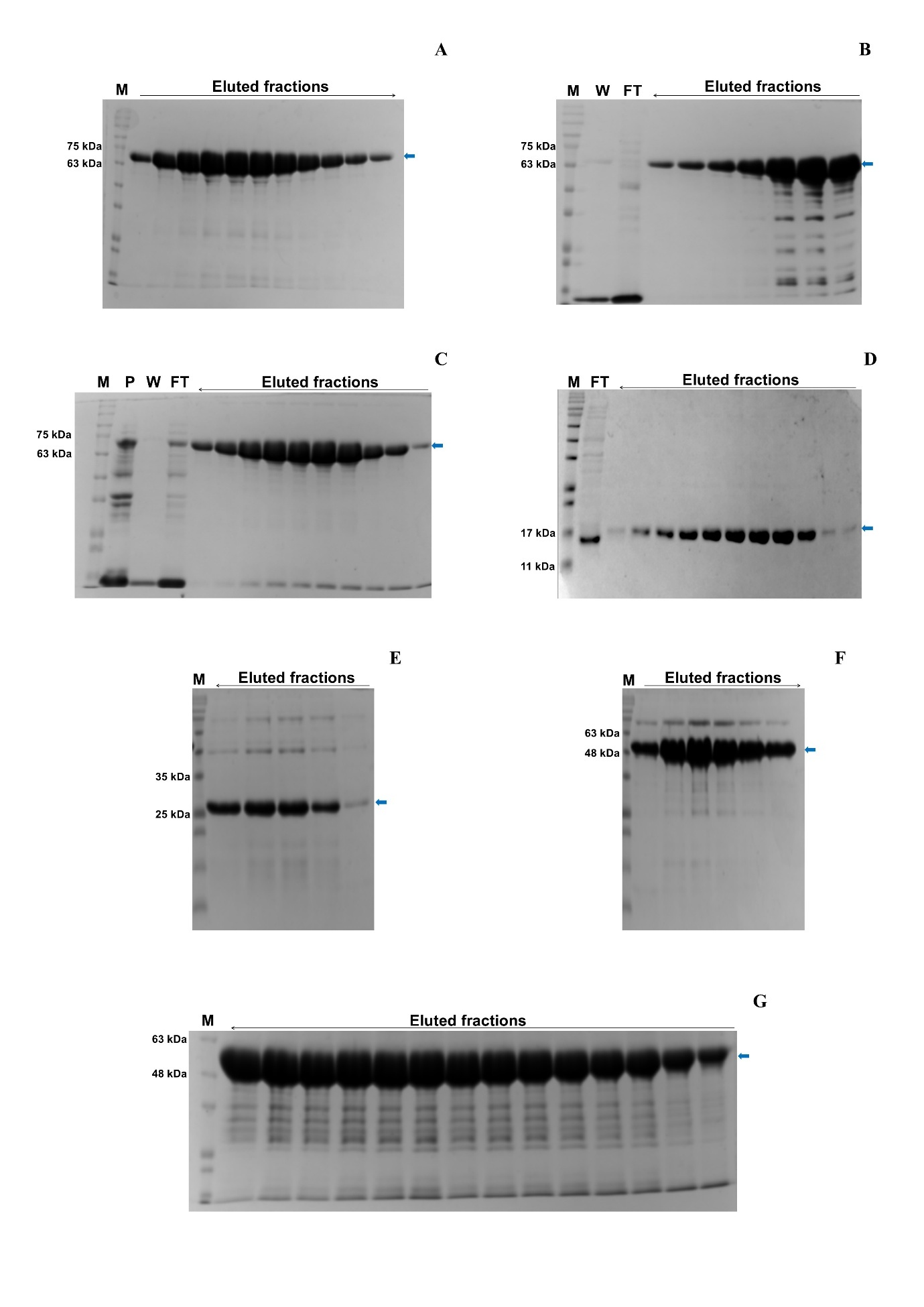

**Fig. S2. Purification of His_6_-tagged human HSF1 and its derivatives used in this study.**

Coomassie-stained 10% SDS-polyacrylamide gel representing the purified HSF1-WT protein (calculated MW ~57 kDa). 20 µl aliquot from each fraction (~200 µl) was loaded into the gel. Arrow on the top indicates the order of fraction collection. The blue arrow indicates the intact HSF1-WT band (A).

Coomassie-stained 10% SDS polyacrylamide gel representing the purified HSF1-ΔLZ1-3. The calculated MW of HSF1-ΔLZ1-3 is ~50 kDa. 20 µl aliquot from each fraction (~200 µl) was loaded into the gel. Arrow on the top indicates the order of fraction collection. The blue arrow indicates the intact HSF1-ΔLZ1-3 band (B).

Coomassie-stained 10% SDS polyacrylamide gel representing the purified HSF1-LZ4m. The calculated MW of HSF1-LZ4m is ~57 kDa. 20 µl aliquot from each fraction (~200 µl) was loaded into the gel. Arrow on the top indicates the order of fraction collection. The blue arrow indicates the intact HSF1-LZ4m band (C).

Coomassie-stained SDS-polyacrylamide gel (15%) representing the purified HSF1-DBD. Calculated MW is ~14 kDa. 20 µl aliquot from each fraction (~200 µl) was loaded into the gel. The arrow on the top indicates the order of fraction collection. The blue arrow indicates the intact HSF1-DBD band (D).

Coomassie-stained SDS-polyacrylamide gel (15%) representing the purified HSF1-ΔRD-LZ4-TAD. The calculated molecular weight of HSF1-ΔRD-LZ4-TAD is ~23.14 kDa. HSF1-ΔRD-LZ4-TAD precipitated out of solution during dialysis, and hence could not be included in the present study. 20 µl aliquot from each fraction (~200 µl) was loaded into the gel. Arrow on the top indicates the order of fraction collection. The blue arrow indicates the intact protein band (E).

Coomassie-stained SDS-polyacrylamide gel (15%) representing the purified HSF1-ΔLZ4-TAD. The calculated molecular weight of HSF1-ΔLZ4-TAD is ~41.81 kDa. 20 µl aliquot from each fraction (~200 µl) was loaded into the gel. Arrow on the top indicates the order of fraction collection. The blue arrow indicates the intact protein band (F).

Coomassie-stained 10% SDS-polyacrylamide gel representing the purified HSF1-ΔTAD. The calculated MW is ~44.6 kDa. 20 µl aliquot from each fraction (~200 µl) was loaded into the gel. The arrow on the top indicates the order of fraction collection. The blue arrow indicates the intact protein band (G).

[M: Protein MW Marker; P: Pellet; FT: Flowthrough; W: Wash]

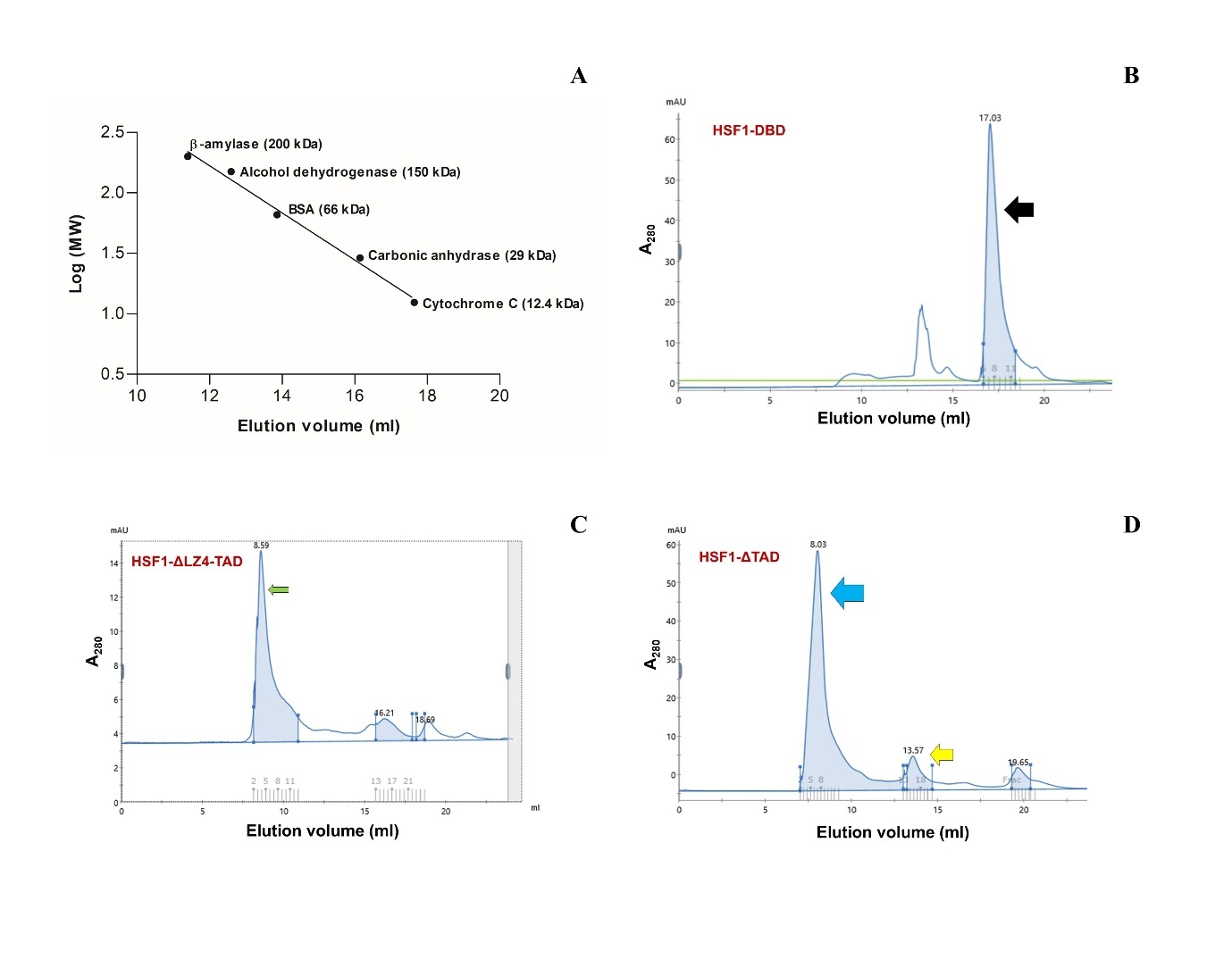

**Fig. S3.** A size exclusion chromatography (SEC) standard curve was obtained by plotting the Log (MW) of the indicated protein standards versus their elution volumes (ml) in a Superdex200 Increase 10/300 GL column. Linear regression analysis was done to obtain the MW of His_6_-tagged HSF1-DBD by putting its elution volume (17.03 ml) in the equation of the straight line (y=-0.1948x+4.5618). The MW obtained from the standard curve agrees well with the calculated MW of the said protein. MWs of other proteins in this study could not be calculated from this curve owing to the presence of intrinsically disordered regions (IDRs) in them (A).

SEC profile of HSF1-DBD. The major peak (eluted at 17.03 ml, indicated by black arrow) corresponds to HSF1-DBD, as suggested by its MW of 17.78 kDa calculated from the SEC standard curve **(Fig. S3A)** by linear regression analysis aligning closely with the actual MW of 16 kDa for his_6_-HSF1-DBD**.** Thus, the major peak was identified as HSF1-DBD, and the shorter peak probably consisted of protein aggregates/ impurities (B).

The SEC profile of HSF1-ΔLZ4-TAD showed a major peak at 8.59 ml (indicated by green arrow), reflecting its constitutive multimeric state, due to the absence of the autoinhibitory LZ4 domain **(Fig. 1A).** The observed elution volume is in good agreement with that of the trimeric/ oligomeric form of HSF1-WT and HSF1-LZ4m **(Fig. 1: C & E)** (C)**.**

The SEC profile of HSF1-ΔTAD revealed a predominance of multimeric species (eluted at 8.03 ml, indicated by blue arrow) with a very small peak of monomeric species (eluting at 13.57 ml, indicated by yellow arrow). We used this derivative directly after the affinity purification for further studies without subjecting it to SEC, as the monomeric species aggregated upon concentration (D).
